# An iPSC-based model of Jacob Syndrome reveals a DNA methylation-independent transcriptional dysregulation shared with X aneuploid cells

**DOI:** 10.1101/2024.06.18.598378

**Authors:** V. Astro, K. Cardona-Londoño, L.V. Cortés-Medina, R. Alghamdi, G. Ramírez-Calderón, F. Kefalas, J. Dilmé-Capó, S. Radío, A. Adamo

## Abstract

Male sex chromosome aneuploidies are frequent genetic aberrations in humans characterized by additional Y or X chromosome complements. Jacob (JS) and Klinefelter syndromes (KS), carrying 47,XYY and 47,XXY chromosomes, respectively, share several clinical features, including sterility, hormonal deficits, neurocognitive delay, and skeletal-muscle defects. Despite the high incidence, a complete understanding of the molecular mechanisms underlying the clinical manifestations in sex aneuploid patients is still elusive. In this study, we generated and characterized the inaugural cohort of 47,XYY human induced pluripotent stem cells (iPSCs). We performed a comprehensive transcriptional analysis, including JS primary fibroblasts, iPSCs, and neural stem cells (NSCs) alongside KS cells. Our study revealed a transcriptional feedback mechanism tuning non-PAR X chromosome genes (NPX) homologs in Y supernumerary cells, a phenomenon not detected in X aneuploid iPSCs. By ectopically modulating the expression of selected NPY genes, we demonstrated a direct transcriptional link between the UTY-UTX gene pair. Furthermore, our analyses identified a shared transcriptomic signature between JS and KS, discernible already at the iPSC stage, with a notable enrichment for processes related to neurological functions. This transcriptomic convergence underscores potential commonalities in the molecular pathways underpinning the pathophysiology of male sex chromosome aneuploidies. Finally, through genome-wide DNA methylation profiling of JS iPSCs, we demonstrated that a supernumerary Y chromosome only minimally impacts the methylation status of 47,XYY cells at the pluripotent stage. Our work reveals critical transcriptional feedback mechanisms and shared gene expression signatures in male sex chromosome aneuploidies, paving the way for a better understanding of their common phenotypic features.

## INTRODUCTION

Jacob Syndrome (JS) is the second most prevalent sex chromosome aneuploidy in males, with a rate of 1:1000 newborns^1–3^. JS is characterized by an additional Y complement in males due to paternal non-meiotic disjunction of the Y chromosome, thus resulting in a 47,XYY karyotype. Despite the high incidence, a comprehensive understanding of the molecular mechanisms underlying the onset of the clinical features of JS patients remains elusive. Frequent phenotypic traits observed in JS patients include tall stature, macrocephaly, macrodontia, clinodactyly, scoliosis, increased testicular volume, and hypotonia^4,5^. Additionally, JS patients commonly exhibit neurological, cognitive, and behavioral deficits, an IQ score lower than siblings, and increased risks of hyperactivity and attention deficits^6–10^. Hormonal deficiencies, including hypogonadism, impaired testicular function, micropenis, and infertility, have also been reported^4,11^. Intriguingly, JS shares several neurocognitive characteristics with Klinefelter Syndrome (KS), such as language disorders, tremors, anxiety, and intellectual disabilities^12^. The shared phenotypic features may stem from the overdosage of genes within the pseudoautosomal regions 1 and 2 (PAR1 and PAR2). The PAR regions are located at the terminal ends of X and Y chromosomes, spanning approximately 2.6 Mb in Xp and Yp (PAR1) and ∼320 Kbp in Xq and Yq (PAR2). Genes within these territories escape X inactivation and are expressed from the three sex chromosomes in 47,XYY (JS) and 47,XXY (KS) cells. The PAR1 region harbors twenty-five genes^13–15^, while PAR2 contains only four genes, with two, *VAMP7* and *SPRY3*, predominantly inactive on the Y chromosome^16,17^. Another potential molecular mechanism underlying the shared neurological features of KS and JS involves the overdosage of non-PAR Y genes (NPY) with homologs on the X chromosome. Of particular relevance could be the overdosage of *NLGN4Y*, *UTY*, *ZFY*, *and DDX3Y* because the non-PAR X (NPX) homologs *NLGN4X, UTX*, ZFX, and *DDX3X*, expressed from both Xs in KS males^18^, have been previously associated with autism spectrum disorder (ASD), intellectual deficits, developmental delay, and behavioral defects^19–26^. Despite the significant impact of sex chromosome aneuploidies in humans and their high incidence among males, few models are available to study the effects of sex gene overdosage on these conditions. Notably, no human induced pluripotent stem cell (hiPSC) models of JS have been described to date, highlighting the urgent need for viable *in vitro* models to untangle the impact of sex chromosome aneuploidies during the earliest stages of human development.

This study recruited three JS patients and generated the inaugural cohort of JS-iPSCs and iPSC-derived neural stem cells (NSCs). We assessed the global transcriptional impact of Y-linked gene overdosage by comparing primary 47,XYY fibroblasts to clonal iPSCs and NSCs. Our data demonstrated the existence of an indirect, compensatory transcriptional mechanism modulating the expression of NPX genes in response to NPY gene overdosage. Moreover, our work couples a cross-tissue transcriptomic and methylomic analysis of supernumerary Y chromosomes and provides innovative insights into Y-mediated modulation of the autosomal genome. Finally, comparing KS and JS primary fibroblast and iPSC transcriptomes, we identified a shared dysregulated signature potentially linked to the common phenotypic outcome of the two diseases. Our work highlights the importance of implementing cellular models of clonal origin with minimized genomic background variability to dig into the molecular mechanisms dysregulated during the embryonic developmental stages of sex chromosomal aneuploidies.

## RESULTS

### 1) Generation of 47,XYY Jacob Syndrome iPSCs and Neural stem cells (NSCs)

We selected a cohort of three non-mosaic 47,XYY JS patient fibroblasts through a bank repository (**Figure 1A**, **Table S1**). We cultured patient fibroblasts under identical conditions and validated the overdosage of expression of the Y-linked genes *UTY*, *KDM5D,* and *ZFY* in JS versus 46,XY cells (**Figure S1A**). We used a virus-free, integration-free somatic cell reprogramming methodology to derive 47,XYY JS-iPSCs (**Figure S1B**). We characterized three independent iPSC clones from each 47,XYY patient (**Figure 1A**) and performed a short-tandem-repeat (STR) analysis to match the genetic profile of the parental fibroblasts and the derived iPSCs (**Table S2**). Next, we assessed the sex chromosome complement of the iPSCs with a dual approach. DNA-FISH assay confirmed the presence of one X and two Y chromosomes in the nuclei of each 47,XYY iPSC clone (**Figure 1B**, **Figure S2**). High-resolution Karyostat assay confirmed a 47,XYY karyotype for all iPSCs generated in this study and excluded the presence of any chromosome rearrangement in all clones except for 1.2 and 2 Mb gain in clones JS1 #B and JS2 #B, respectively (**Figure 1C**, **Figure S3**). Moreover, all iPSC lines exhibited canonical pluripotent morphology and expressed high levels of the pluripotency markers *NANOG*, *SOX2,* and *OCT4* at the mRNA level (**Figure S4A**) and of the NANOG, SOX2, and SSEA4 proteins by immunofluorescence staining (**Figure S4B-C**). To model the consequences of Y aneuploidy during the earliest stages of human neurodevelopment, we differentiated 47,XYY and 46,XY iPSCs into neural stem cells (NSCs), self-renewing multipotent intermediates from which the major cell types of the adult central nervous system arise^27^. JS and 46,XY iPSCs were differentiated into NSCs *in vitro* through a stepwise differentiation protocol (**Figure S5A**), leading to the robust induction of the neuroepithelial precursor marker *NES* and the neuronal marker *TUBB3* accompanied by the concomitant shut-down of the pluripotency gene *OCT4* (**Figure S5B-D**).

**Figure 1:**
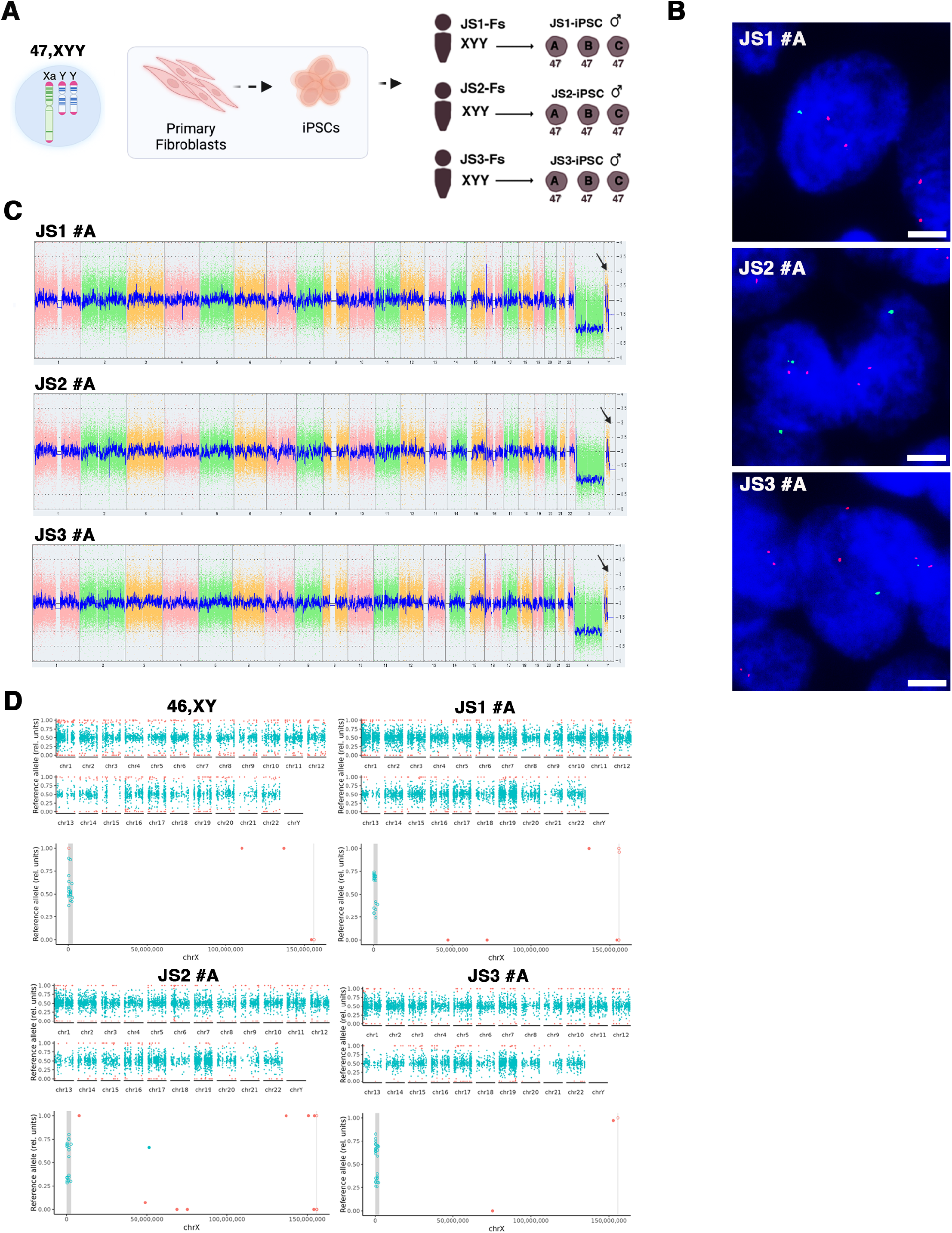
Characterization of 47,XYY JS iPSCs. **A)** Schematic of the nine 47,XYY iPSC clones generated in this study. **B)** Representative DNA-FISH images of X chromosome (green) and Y chromosome (red) in JS-iPSCs. DNA was stained with DAPI (Blue). Scale bar = 5 µm. **C)** KaryoStat+ whole genome view. The whole genome view displays all somatic and sex chromosomes in one frame with a high-level copy number. The smooth signal plot (right y-axis) is the smoothing of the log2 ratios, which depict the signal intensities of probes on the microarray. The pink, green, and yellow colors indicate the raw signal for each chromosome probe, while the blue represents the normalized probe signal, which is used to identify copy numbers and aberrations. Black arrows indicate the gain of an additional Y chromosome. **D)** Scatter plot profiles of coupled WES analysis and allele-specific RNA-Seq analysis performed on autosomal (upper panel) and sex chromosomes. The X chromosomes are highlighted in the lower panel. Plots show the mono-(orange dots) or biallelic (light blue dots) gene expression status in two representative karyotypes (46,XY and 47,XYY). Gray rectangles indicate PAR1 and PAR2 regions, respectively. Solid dots indicate non-PAR genes; open dots show PAR genes. The scatter plot of iPSC clone JS2#A assigns a biallelic SNP to NUDT10. See the Material and Methods section.

### 2) Supernumerary Y chromosome increases PAR and NPY gene expression levels

Genes within the human PAR regions on the Y and the X are expressed from both sex chromosomes in male and female cells^28^. We previously demonstrated that, in the presence of one or more supernumerary X chromosomes, the expression of PAR1 genes mirrors the number of Xs^29^. Thus, to ascertain whether a similar gene overexpression would be detected in supernumerary Y cells and to verify the biallelic expression profile of the PAR genes in JS, we coupled transcriptomic analysis to whole exome sequencing (WES) and allele-specific expression (ASE) analysis in 47,XYY fibroblasts, iPSCs, and iPSCs-derived NSCs. As expected, most autosomal genes displayed a biallelic expression profile. On the other hand, biallelically expressed SNPs were detected exclusively in the PAR1 region on the X and Y chromosomes (**Figure 1D**, **Figure S6**). Notably, given the bioinformatic convention of assigning PAR regions on the X while masking the Y chromosome^30^, every expressed PAR SNP is automatically mapped on chromosome X. Moreover, since the supernumerary Y chromosome in 47,XYY cells originates from a non-disjunction of sister chromatids of paternal origin^31^ there are no biallelic expressed SNPs assigned to the Y chromosome (**Figure 1D**, **Figure S6**). Interestingly, biallelic expressed SNPs mapping to the PAR1 region in JS samples were distributed around the values 30% and 60%, suggesting that the X and the two identical Y chromosomes contribute similarly to the total expression level of the PAR1 transcripts. Coherently, the expressed SNPs mapping to the PAR1 region in 46,XY controls clustered around 50% (**Figure 1D**).

Next, we investigated the impact of the supernumerary Y chromosome on the global transcriptome by profiling fibroblasts, iPSCs and NSCs obtained from 47,XYY and 46,XY individuals (**Figure 2A**). The unsupervised principal component analysis (PCA) of 47,XYY and 46,XY transcriptomes revealed that samples cluster according to cell type and karyotype (**Figure 2B**). We detected 13153, 14900, and 13989 expressed genes in fibroblasts, iPSCs, and NSCs, respectively (**Table S3**). First, we analyzed the expression of Y and X-linked genes residing within different chromosomal territories, the PAR, and the NPY and NPX regions. The Y chromosome harbors a restricted number of genes, most of which have a homolog on the X (**Figure 2C**). We counted 14, 18, and 16 NPY-expressed genes in fibroblasts, iPSCs, and NSCs, respectively (**Figure 2D-F**, **Table S3**). The expressed X-linked genes in fibroblasts, iPSCs, and NSCs are 439, 545, and 515, respectively. By studying the expression profile of PAR1 and PAR2 genes in the different cell types, we discovered that PAR gene expression is differentially regulated in a tissue-specific fashion (**Figure S7**).

**Figure 2:**
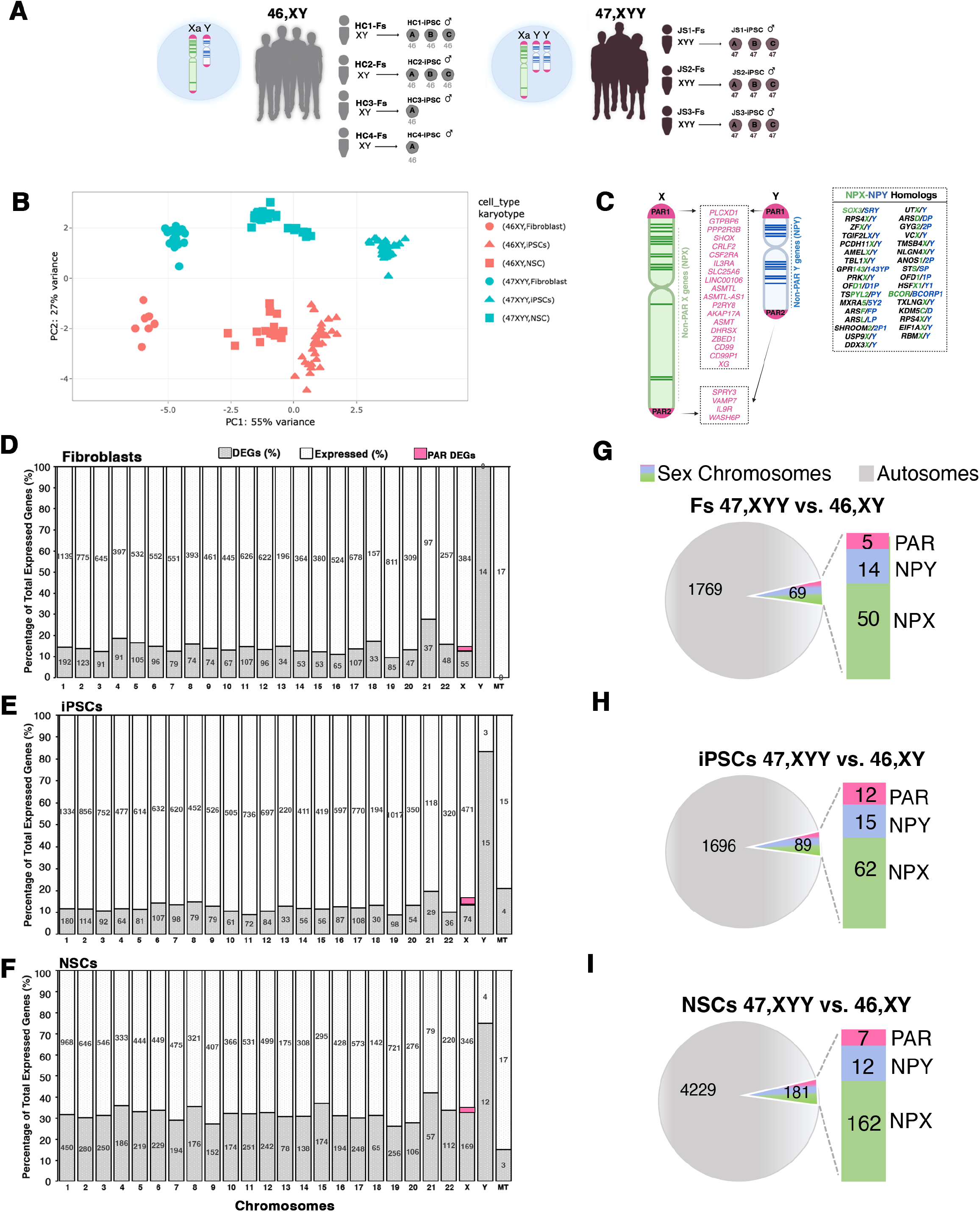
Transcriptional impact of Y chromosome aneuploidy. **A)** Illustration of the 47,XYY patient and control 46,XY fibroblasts (Fs) and derived-iPSC clones used in the study. Three 47,XXY fibroblast lines have been reprogrammed to generate three independent 47,XYY iPSC clones each. HC, Healthy controls; JS, Jacob Syndrome. The control 46,XY Fibroblasts, and iPSCs have been described in *Astro et al*., *2021*^29^ and *Astro et al*., *2023*^33^. **B)** PCA analysis of all samples used in the study. 47,XYY, blue symbols; 46,XY, orange symbols **C)** Schematic of sex human sex chromosomes listing the genes encoded from the X and Y chromosome shared pseudoautosomal territories, PAR1 and PAR2, and from the divergent NPX and NPY regions. **D-F)** Bar plots of the percentage of differential expressed genes (DEGs) over the total number of expressed genes in fibroblast, iPSCs, and neural stem cells (NSCs). The specific number of DEGs and expressed genes are shown inside the bars for each chromosome. PAR DEGs are indicated in pink and assigned only to the X chromosome. MT, mitochondrial genes. **G-I)** Pie chart showing the number of autosomal and sex chromosome DEGs in each cell type. The number of X and Y-linked PAR, NPX, and NPY DEGs are indicated in the bars. Log_2_FC >|0.58|, FDR <0.05. Fs, Fibroblasts.

The differential expression analysis (DEA) performed on 47,XYY and 46,XY karyotypes identified 1838, 1789, and 4413 differentially expressed genes (DEGs) in fibroblasts, iPSCs and NSCs, respectively (FDR<0.05 and Log_2_FC >|0.58|) (**Figure 2D-F**, **Figure S8, Table S4**). Considering only the NPY-expressed genes, 14 (93%) in fibroblasts, 15 (83%) in iPSCs and 12 (75%) in NSCs were identified as DEGs (**Figure 2D-I**). Importantly, all NPY DEGs were upregulated in the three cell types except for *MXRA5Y,* which was downregulated in fibroblasts (**Figure S9A-C)**. All PAR DEGs were upregulated and localized in the PAR1 region. *WASH6P* was the only PAR2 gene differentially expressed in iPSCs and NSCs (**Figure 2G-I, Figure S9D-F**). Among the NPY DEGs, ten are shared across all cell types, and all except *TTTY14* have NPX homologs (**Figure 3A**). On the contrary, the identity of differentially expressed PAR genes varies across cell types (**Figure 3B**, **Figure S9D-F**). Notably, a correlation analysis of the logarithmic, base 2, fold change of NPY and PAR DEGs comparing fibroblasts, iPSCs, and NSCs revealed a comparable magnitude of dysregulation of NPY and PAR genes across 47, XYY cells, except for NPY genes in NSCs against iPSCs (**Figure 3C-D**).

**Figure 3:**
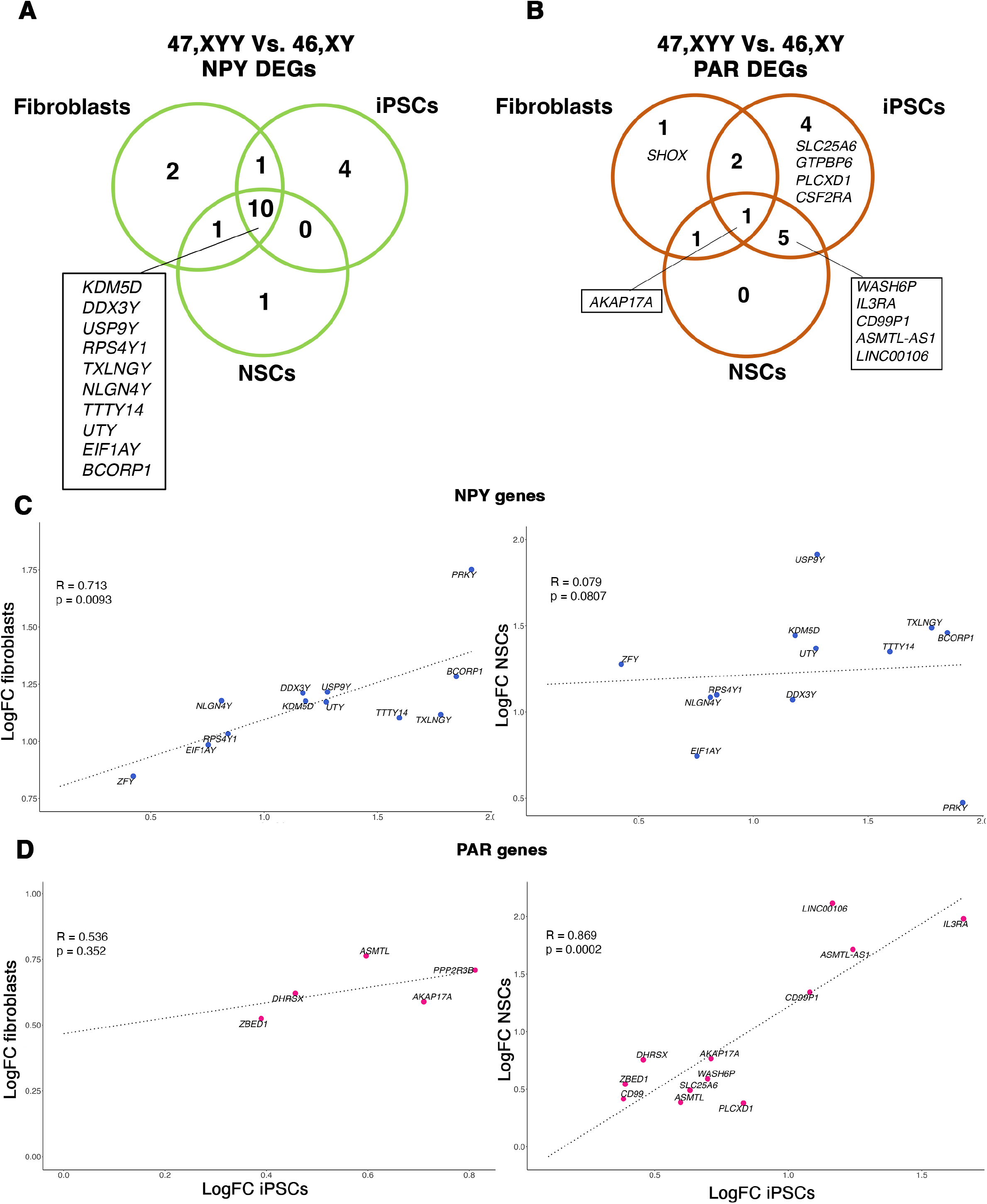
Transcriptomic profiling of sex chromosome gene expression in 47,XYY cells. Venn diagrams of shared **A**) NPY DEGs and **B**) PAR DEGs in 47,XYY vs. 46,XY fibroblasts, iPSCs, and NSCs. C-D) Scatterplot of the Log2 Fold Change (Log2FC) comparison for NPY genes (**C**) and X- and Y-linked PARs(D) in fibroblasts vs. iPSCs (left) and IPSCs vs. NSCs (right). FDR < 0.01. NPY genes are indicated with blue dots. PAR genes are pink. The Pearson correlation coefficient of the regression line (R) and p values are shown.

### 3) NPY gene overdosage leads to a compensatory transcriptional downregulation of NPX homolog genes

Next, we investigated whether the presence of a supernumerary Y chromosome would lead to a transcriptional compensatory modulation of NPX genes. The DEA identified the X-linked genes whose expression positively or negatively changes in response to Y gene overdosage in fibroblasts, iPSCs, and NSCs **(Figure S10)**. Notably, a subclass of NPX genes has an NPY homolog for which the degree of functional and structural similarities is mainly unexplored (**Figure 2C**). Thus, we compared the expression of NPX and NPY homologs and identified *NLGN4X, EIF1AX, DDX3X, and TXLNGX as* genes significantly downregulated in 47,XYY iPSCs and NSCs. Conversely, their Y-homologs *NLGN4Y, EIF1AY, DDX3Y, and TXLNGY* were upregulated in both 47,XYY cell types (FDR < 0.01) (**Figure 4A-B**). The NPX genes *ZFX*, *ANOS1*, and *BCOR* displayed an opposite expression trend compared to their NPY homologs in iPSCs but not NSCs. Intriguingly, the histone demethylase *UTX/KDM6A* and *USPX* follow the same expression trend of their Y homologs (*UTY* and *USPY*) and are upregulated in iPSCs and NSCs. *KDM5C* is the only NPX gene consistently insensitive to increased Y paralogous (*KDM5D*) expression in both cell types (**Figure 4A-B**). Next, we assessed the correlation between the Y chromosome dosage and the expression of sex chromosome genes. To this aim, we generated a linear regression model with two (47,XYY), one (46,XY), or zero (46,XX) Y chromosome complements by integrating the 46,XX iPSC transcriptomes previously obtained in our laboratory^29,32,33^ (**Table S5**). Our results demonstrate that a subset of NPX genes undergoes an inversely transcriptional regulation proportional to the number of Y chromosomes (**Figure 4C**). To test the intriguing hypothesis that NPY genes directly modulate the expression of their X homologs, we acutely knocked down *ZFY*, *NLGN4Y, DDX3Y, and UTY* in 47,XYY iPSCs using siRNAs (**Figure 4D**). The knockdown efficiently reduced the expression of the targeted genes in three independent 47,XYY iPSCs (**Figure 4E**). Importantly, the *ZFX*, *NLGN4X,* and *DDX3X* expression was insensitive to *ZFY*, *NLGN4Y,* and *DDX3Y* knockdown. On the other hand, upon *UTY* knockdown, the expression of *UTX* was significantly downregulated (**Figure 4E**). Our results suggest a mechanism of transcriptional modulation of NPX homologs in JS iPSCs and NSCs, although a direct reciprocal regulation of single NPY-NPX gene pairs could only be detected for *UTY*-*UTX* in our model system.

**Figure 4:**
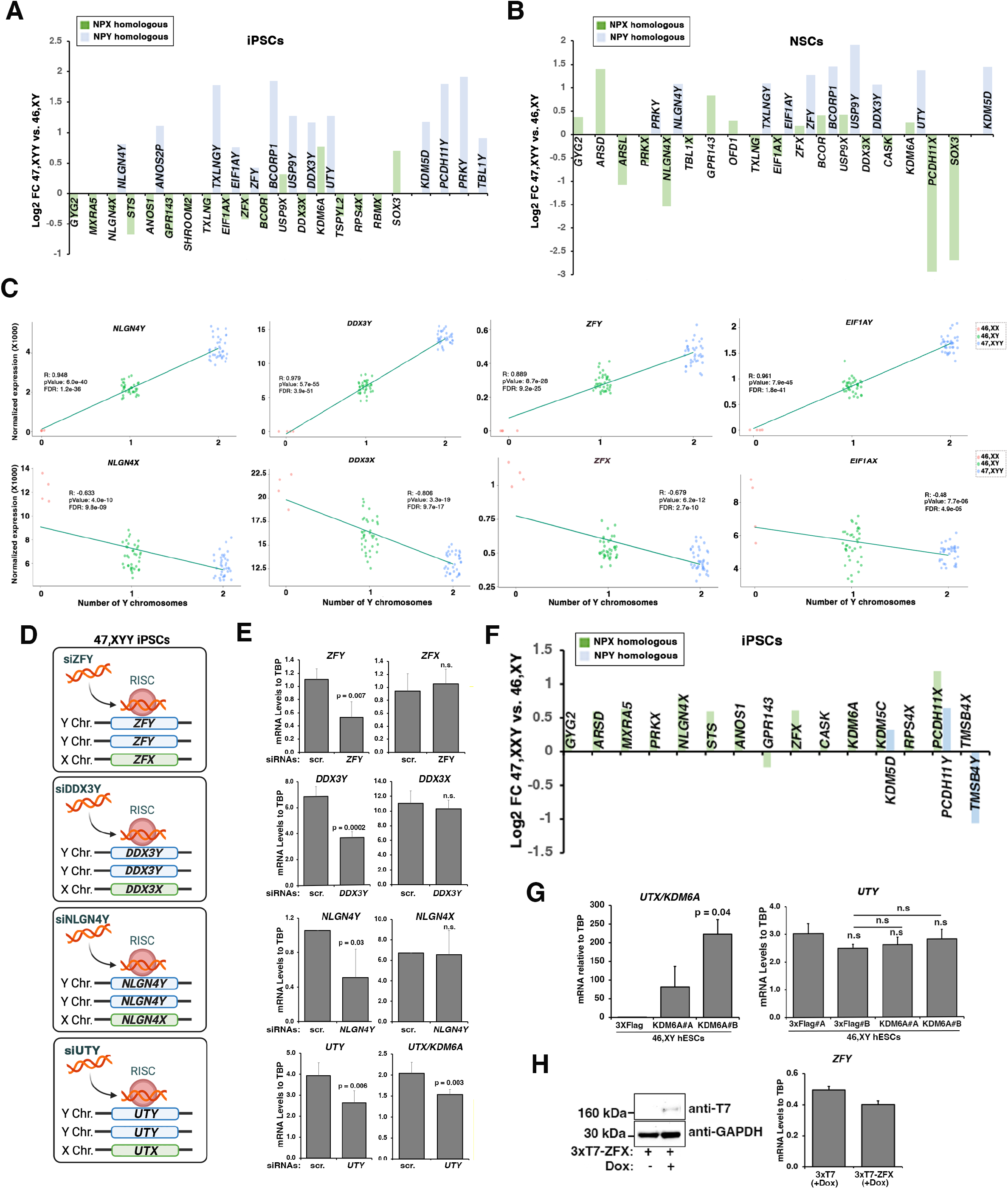
NPY genes modulate the expression of their homologs on the X chromosome. **A-B)** Log2 fold change of NPY (light blue) and NPX (green) expression in the contrast 47,XYY versus 46,XY in (A) iPSCs and (B) NSCs. DEGs with FDR < 0.01 are plotted. **C)** Linear regression analysis of TMM normalized gene expression in iPSCs carrying zero (46,XX), one (46,XY), or two (47,XYY) Y chromosomes. Pearson correlation coefficients (R), p values, and FDR are indicated. Top: NPY genes show a positive expression correlation with Y number. Bottom: NPX homologs show an opposite correlation trend. **D)** Schematic of siRNA experiments. 47,XYY iPSCs have been transfected with siRNA against *ZFY*, *DDX3Y*, *NLGN4Y*, and *UTY*. **E)** Taqman assays are used to analyze the expression of the indicated mRNAs in control cells transfected with either siRNA scramble (scr.) or siRNAs specific to the indicated targets. Bars represent the average ± std of four independent experiments on iPSC clones derived from the three patients. Student’s t-test, one-tailed distribution, two-sample unequal variance. P values are indicated. **F)** Log2 fold change of NPX (green) and NPY (light blue) expression in the contrast 47,XXY versus 46,XY in iPSCs. DEGs with FDR < 0.01 are plotted. **G)** Gene expression by Real-time qPCR for *KDM6A* (left) and Taqman assay for *UTY* (right) in 46,XY hESCs stably integrating either empty 3X-Flag (control) or 3X-FlagKDM6A vectors. Two independent hESC clones (KDM6A#A and KDM6A#B) have been tested. Bars are the average ± std of three independent experiments. Student’s t-test, two-tailed distribution, two-sample unequal variance. P values are indicated. The significance is set at p values < 0.05. **H)** Left: Western Blot showing the inducible overexpression of ZFX protein in hESCs H1 treated with 1 ug/ml Doxycycline (+Dox) or not treated (-Dox). Right: Taqman assay assessing ZFY expression levels in Doxycycline (+Dox) treated hESCs. n.s = not significant.

Next, we investigated if, similarly to JS, the number of supernumerary X chromosomes in KS iPSCs would correlate with the downregulation of the NPY genes. Integrating a dataset of 47,XXY and 46,XY iPSCs transcriptomes^29,33^, we discovered that none of the NPY genes was significantly downregulated in KS iPSCs, except for *TMSB4Y* (**Figure 4F**). We concluded that NPY genes are mainly insensitive to X dosage. To further validate this evidence, we engineered H1 human embryonic stem cells (46,XY hESCs) to overexpress *ZFX* or *UTX* stably, and we did not observe significant changes in *ZFY* and *UTY* levels (**Figure 4G-H**). Our data indicate that Y chromosome overdosage leads to NPX gene expression modulation but not *vice versa*, thus suggesting a unidirectional regulatory mechanism of Y genes on their X homologs.

### 4) Y chromosome aneuploidy leads to global transcriptional dysregulations

Next, we investigated the impact of Y aneuploidy on the global transcriptome of fibroblasts, iPSCs, and NSCs. We performed a gene ontology (GO) analysis, selecting genes commonly dysregulated in at least two cell types (**Figure 5A**). Remarkably, we identified enrichment for terms related to neuron system development, axon guidance, cell migration, and neuron projection (**Figure 5B**), all processes particularly significant for JS phenotypic features. We narrowed our analysis to iPSC DEGs responsive to Y chromosome dosage. To this end, we built a correlation model using iPSCs with zero (46,XX), one (46,XY), and two (47,XYY) Y chromosomes, and we detected 854 genes sensitive to Y-dosage. Additionally, we crossed Y dosage-sensitive genes with the DEGs in iPSCs (47,XYY vs. 46,XY) and identified 622 DEGs (FDR <0.01) whose expression changes proportionally to the number of Y chromosomes (**Figure 5C**). The GO analysis on these DEGs confirmed that genes associated with neuronal morphogenesis and function are dysregulated at the pluripotent stage and sensitive to Y chromosome dosage (**Figure 5D-E**). Among the Y-sensitive autosomal genes, *NBPF8* and *SLIT1* had the highest positive correlation, while *PIK3R1* and *PTPN9* had the highest negative correlation with the number of Ys (**Figure 5F).** *NBPF8* and *SLIT1* are both involved in nervous system development. *NBPF8* is a member of the neuroblastoma breakpoint family (NBPF) and is characterized by tandemly repeated copies of DUF1220 protein domains. Gene copy number variations of these domains have been associated with developmental and neurogenetic diseases characterized by human brain size defects and cognitive disability^34^. *SLIT1* functions include axon extension and negative chemotaxis^35,36^. *PIK3R1* plays a crucial role in insulin metabolism by modulating glucose uptake and glycogen synthesis in insulin-sensitive tissues^37^. *PTPN9* is a protein tyrosine phosphatase (PTP) family member whose haploinsufficiency is associated with vascular, bone, and neural tube abnormalities in mice^38^.

**Figure 5:**
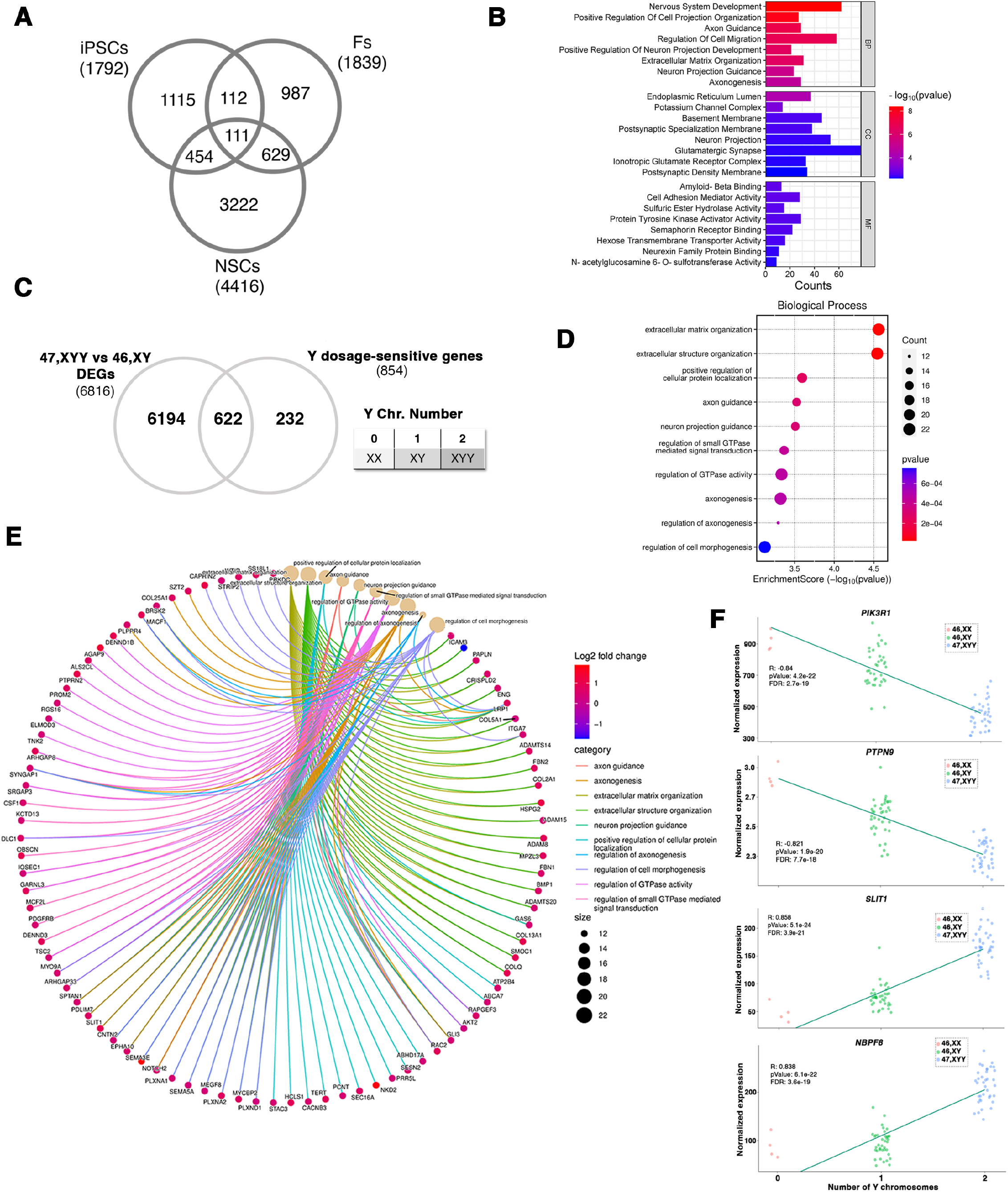
Autosomal response to Y chromosome aneuploidy. **A)** Venn diagram of shared DEGs comparing 47,XYY vs. 46,XY, across three cell types. **B**) GO analysis of 1306 DEGs shared in at least two comparisons. **C**) Venn diagram of DEGs in the 47,XYY vs 46,XY contrast and sensitive to Y chromosome dosage. Dosage-sensitive genes are listed in Table S5. FDR < 0.01 and Pearson correlation value > |0.48|. D) Gene ontology analysis on the DEGs responsive to Y chromosome number showing significantly enriched biological processes. E) Network analysis plot of genes belonging to enriched biological processes of the GO analysis shown in (**D**). F) Linear regression analysis of the TMM normalized gene expression in iPSCs carrying zero (46,XX), one (46,XY), or two (47,XYY) Y chromosomes. The four top hits of autosomal genes with dosage sensitivity to the Y number are shown. Pearson correlation coefficients (R), P values, and FDR are indicated.

### 5) The presence of an extra Y chromosome results in subtle changes in DNA methylation

To understand the impact of an extra Y chromosome on global DNA methylation, we performed a genome-wide methylation analysis (GWMA) using reduced representation bisulfite sequencing (RRBS-Seq) on primary fibroblasts and iPSCs. The PCA of the general distribution of CpGs in 47,XYY and 46,XY demonstrated that the global methylation is primarily influenced by cell type rather than karyotype (**Figure S11A**). We found that the number of significant differentially methylated CpGs in fibroblasts is higher than in iPSCs (**Figure 6A-B**). Next, we assessed the methylation status of the Y chromosome in the two cell types. Primary fibroblasts displayed a heterogenous pattern of CpG methylation, potentially ascribable to the non-clonal nature of this cell population (**Figure 6C**)^39^. On the other hand, methylated CpGs in clonal iPSC lines are significantly more homogenous (**Figure 6D**). Finally, we analyzed the differentially methylated regions (DMRs) comparing 47,XYY and 46,XY cells and identified 4197 and 63 DMRs in fibroblasts and iPSCs, respectively (**Figure S11B, Table S6**). Overall, the small number of DMRs identified in 47,XYY versus 46,XY iPSCs indicates that the supernumerary Y minimally affects the global DNA methylation landscape at this developmental stage. On the other hand, we observed a significant hypermethylation trend in 47,XYY primary fibroblasts (**Figure 6E**). In these cells, the DMRs in autosomal chromosomes represent ∼ 97% of the total detected DMRs (**Figure S11B**) and are uniformly distributed among intergenic, promoter, and gene body genomic regions (**Figure 6F**). The DMRs detected in sex chromosomes (3%) are also predominantly hypermethylated and mostly intergenic. Interestingly, we identified only a few hypermethylated DMRs (8) on the Y chromosome (**Figure 6G**).

**Figure 6:**
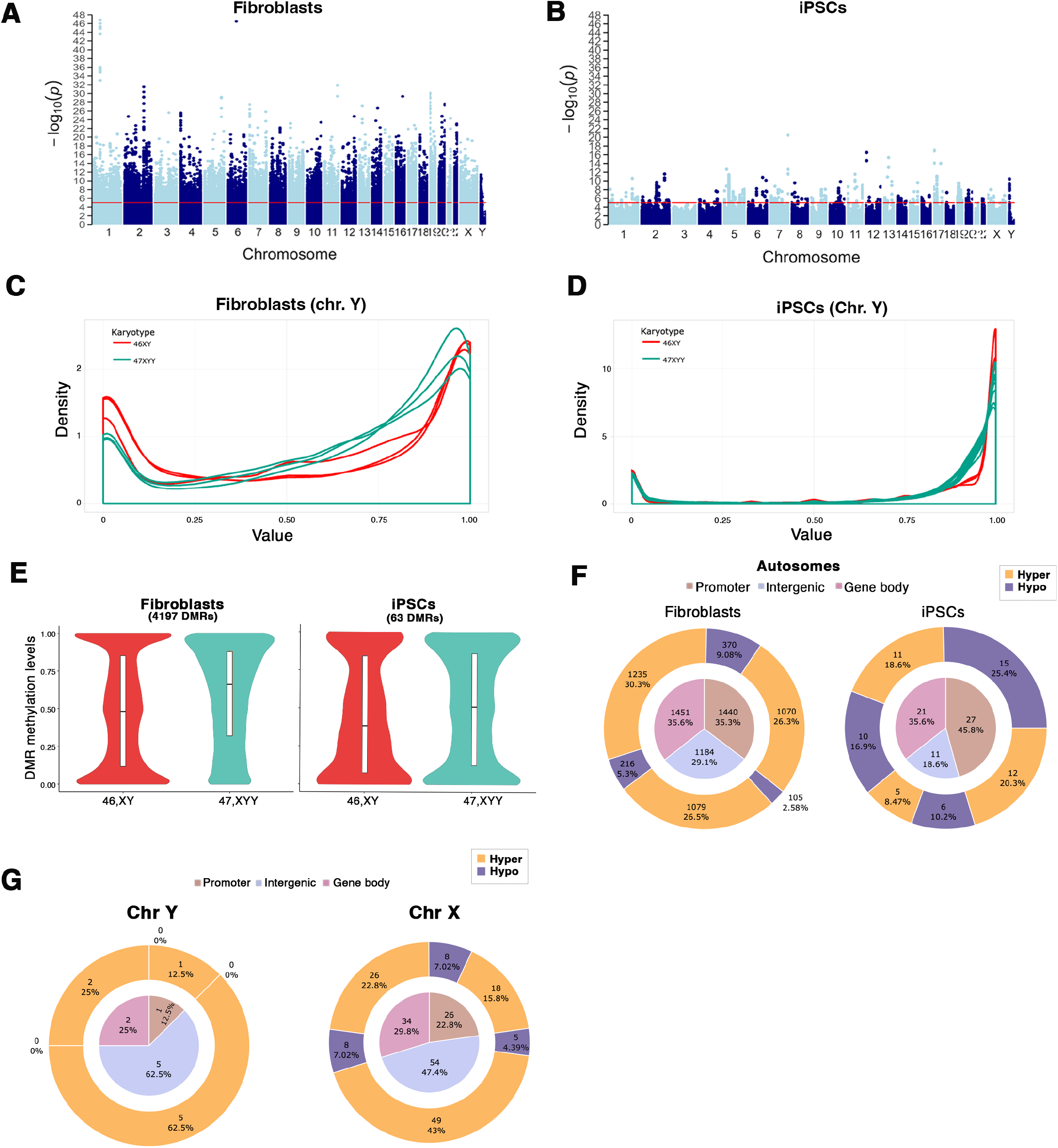
DNA Methylation profile of 47,XYY and 46,XY in fibroblasts and iPSCs. **A-B)** Manhattan Plots showing the methylation distribution in CpG sites in 47,XYY and 46,XY in A) fibroblasts and B) iPSCs. **C-D)** Density plots of the CpG methylation ratios in Y chromosome in 47,XXY vs. 46,XY in fibroblasts (C) and (D) iPSCs. **E)** Violin distribution plot in DMR methylation levels in 47XYY vs. 46XY fibroblasts and iPSCs. The X-axis represents the comparison karyotype, and the Y-axis represents the methylation level value. The inner Boxplot represents the ends of the first and third quartile, and the central slash represents the median. **F)** Nested pie chart of the distribution of autosomal DMRs in Fibroblast (left) and iPSCs (right). **G)** Nested pie chart of the distribution of DMRs in Y (left) and X (right) chromosomes in fibroblasts. The Inner pie represents the proportion of DMR located in the gene promoter (brown), gene body (pink), and intergenic regions (blue). The Outer donut represents the proportion of the DMRs in the inner pie categories that are hypermethylated (orange) or hypomethylated (blue).

### 6) Identification of a shared transcriptional signature in Y and X chromosome aneuploidies

Given JS and KS patients’ highly similar phenotypic features, we investigated if a shared transcriptional impact of Y and X overdosage in JS and KS fibroblasts and iPSCs could be identified. We used a cohort of eight primary KS patients’ fibroblasts and 18 iPSC clones 47,XXY derived in our previous studies^29,33^ to perform a comparative analysis on 47,XYY and 47,XXY cells (**Figure 7A**). First, we determined the impact of a supernumerary X or Y chromosome on X-linked genes by plotting the X-linked gene density distribution in 46,XX, 47,XXY, and 47,XYY normalized on 46,XY iPSCs (**Figure 7B)**. We found that 47,XYY and 47,XXY have a similar density trend of increased gene expression along the terminal Xp arm, including but not limited to PAR1. Also, our findings highlight that JS-iPSCs express PAR genes at a higher level than KS cells. Moreover, while 47,XXY and 46,XX have a similar expression density on the Xq arm, indicating that the second X leads to a similar transcriptional contribution in female and male cells, the additional Y chromosome leads to a mild downregulation along the X chromosome, thus diverging from both KS and female cells (**Figure 7B)**. Next, we compared the contrasts 47,XYY versus 46,XY and 47,XXY versus 46,XY in fibroblasts and iPSCs and identified 30% and 31% commonly dysregulated X-linked DEGs, respectively (**Figure 7C-D, Table S7)**. Importantly, half of the shared DEGs are PAR genes, and they are all upregulated versus 46,XY cells (**Figure 7E).** In iPSCs, the 47,XYY vs. 46,XY contrast displays a higher fold change increase of the shared PAR DEGs compared to 47,XXY vs. 46,XY contrast, thus corroborating the hypothesis that supernumerary Y and X chromosomes contribute differently to PAR gene expression levels (**Figure 7E-F**). However, while clonal cell populations such as iPSCs allow a more quantitative evaluation of the transcriptional contribution of an extra Y or X chromosome, primary cells are more subjected to confounding effects due to cell heterogeneity (**Figure S12**). Finally, we performed an interaction analysis to identify the dysregulation trends among the shared DEGs in 47,XYY and 47,XXY. In fibroblasts, out of 646 common DEGs, 80% are concordantly upregulated (331) or downregulated (190) (**Figure 8A-B, Table S7**). The GO analysis performed on the concordant DEGs highlighted terms related to KS and JS clinical features, such as urogenital system development, neuron projection development, and synapse organization (**Figure 8C-D**). In iPSCs, out of 246 common DEGs, 74% are concordantly upregulated (140) or downregulated (40) (**Figure 8E-F**) and significantly enriched for multiple GO terms related to voltage-gated ion channel activities and immune response (**Figure 8G**). Collectively, our results suggest that a shared transcriptomic signature associated with sex chromosome aneuploidies in males could constitute the molecular landscape, leading to the common phenotypic features in JS and KS patients.

**Figure 7:**
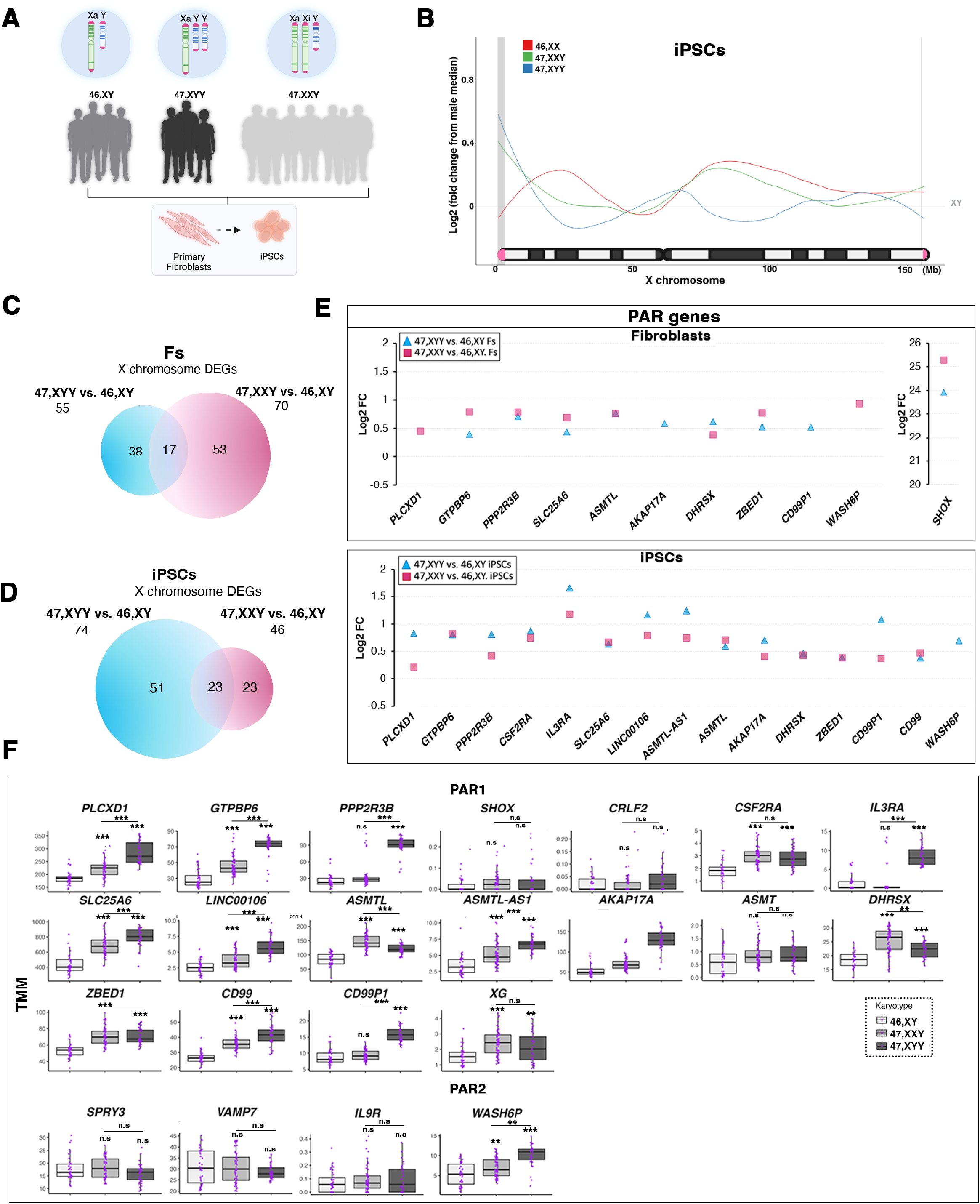
Y and X chromosome aneuploidies contribute differently to PAR gene overdosage. **A)** Schematic of 46,XY, 47,XYY, and 47,XXY fibroblasts and iPSC cohort analyzed in the present study. Fibroblasts from 4 males, 3 JS, and 8 KS patients have been used to derive eight 46,XY, 9 47,XYY, and 20 47,XXY independent iPSC clones. **B)** Moving average line plot (loess fit, span 0.45) along the X chromosome showing the Log_2_ fold change from control 46,XY iPSCs. Gray vertical boxes indicate PAR1 and PAR2 regions. The gray horizontal line represents no theoretical deviations from control 46,XY samples. **C-D)** Venn diagram showing overlapping X-linked DEGs in the 47,XYY versus 46,XY (Blue circle) and 47,XXY versus 46,XY (Pink circle) contrasts in (C) Fibroblasts and (D) iPSCs. FDR<0.05 and LogFC>|0.58|. **E)** Log_2_ fold change of detected PAR genes in 47,XYY and 47,XXY versus controls, fibroblasts (top), and iPSCs (bottom). **F)** Box plot of TMM normalized PAR gene expression in 46,XY, 47,XXY, and 47,XYY iPSCs. Each Purple dot represents an independent RNA sample. The significance of the comparison between 47,XXY and 47,XYY versus 46,XY was calculated using the one-way ANOVA and pairwise comparison with Post Hoc Tukey HSD. **p < 0.01; ***p < 0.001. n.s = not significant.

**Figure 8:**
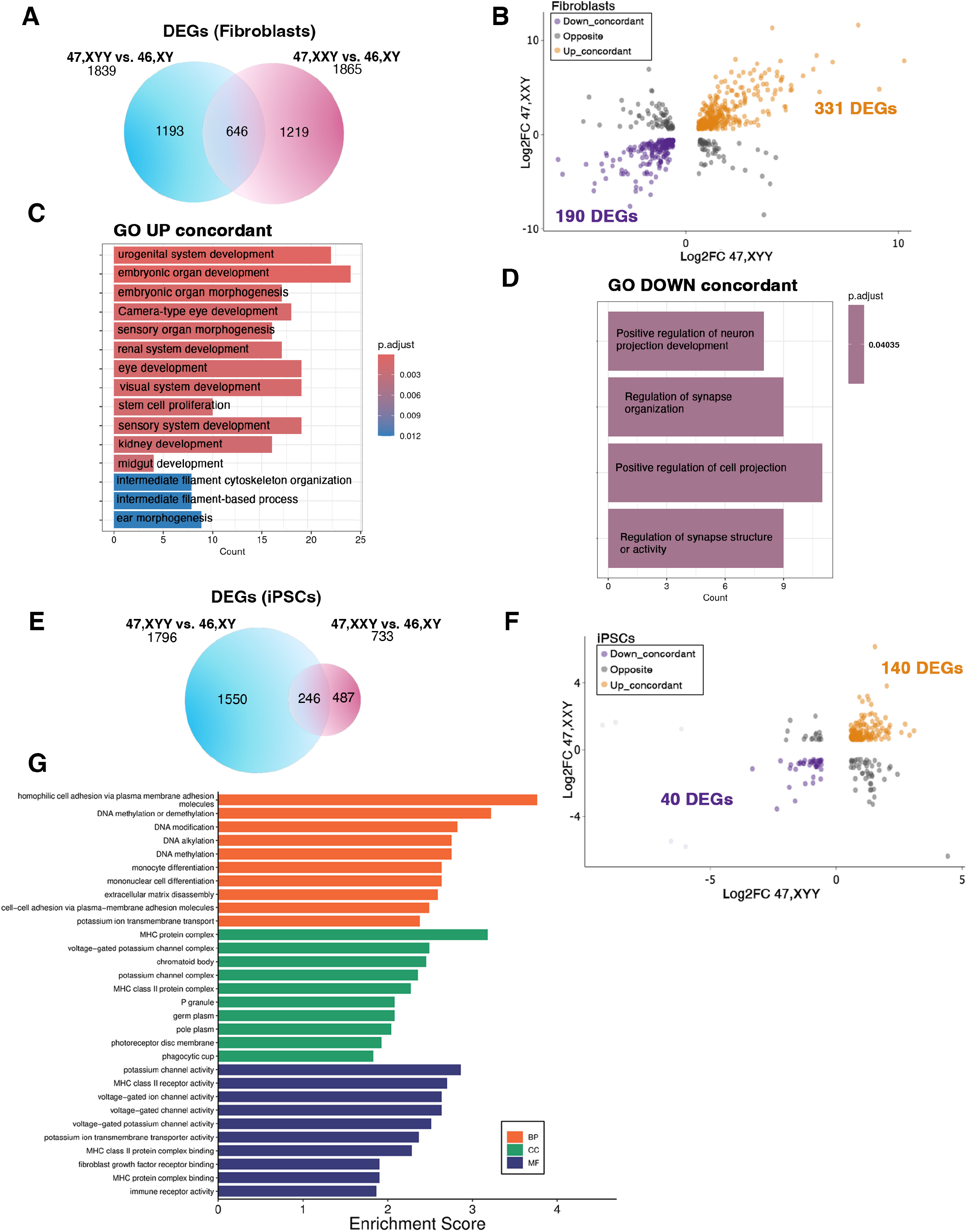
The shared dysregulated genes in X and Y aneuploidies define the transcriptomic signature of sex aneuploidies. **A)** Venn diagram showing overlapping DEGs in the 47,XYY versus 46,XY (Blue circle) and 47,XXY versus 46,XY (Pink circle) contrasts in Fibroblasts. FDR<0.05 and LogFC>|0.58|. **B)** Genome-wide interaction analysis showing the number of DEGs that display an opposite or concordant dysregulation trend in the two contrasts in fibroblasts. See **Table S7** for the complete gene list. **C-D)** Gene ontology enrichment analysis on concordantly (C) upregulated or (D) downregulated DEGs in the KS and JS versus 46,XY fibroblasts. **E)** Venn diagram showing overlapping DEGs in the 47,XYY versus 46,XY (Blue circle) and 47,XXY versus 46,XY (Pink circle) contrasts in iPSCs. **F)** Genome-wide interaction analysis on DEGs with opposite or concordant dysregulation in the two contrasts in iPSCs. See **Table S7** for the complete gene list. **G)** Gene ontology enrichment analysis on the shared concordant DEGs in the KS and JS versus 46,XY fibroblasts. FDR<0.05 and LogFC>|0.58|. BP, Biological processes, MF, Molecular Functions, CC, cellular components.

## DISCUSSION

Our study delves into the intricate biological implications of an additional Y chromosome in Jacob Syndrome (JS), drawing parallels to X aneuploidies in males. Through the successful generation and meticulous characterization of 47,XYY induced pluripotent stem cells (iPSCs) from non-mosaic JS patient fibroblasts, we inaugurated a pivotal understanding of the effects of Y chromosome aneuploidy in human pluripotency and its subsequent impact during differentiation towards disease-relevant lineages. Notably, our methodology hinges upon an mRNA-mediated reprogramming approach, a technique validated for its efficacy in mitigating reprogramming-induced spontaneous sex chromosomal loss, interclone heterogeneity, and insertional mutagenesis events^29,40–43^. This approach ensured the establishment of a homogeneous cellular platform, thereby furnishing a conducive milieu for *in vitro* studies.

Prior investigations into the transcriptional impact of Y and X chromosome aneuploidies predominantly relied on primary patient cells or immortalized lymphoblastoid cell lines^44–52^. While these studies have been instrumental in unraveling common cellular dysregulations associated with these conditions across diverse patient cohorts, they are inevitably encumbered by intrinsic heterogeneity stemming from different genetic backgrounds, ethnicity, age, environmental factors, and potential cryptic mosaicism across various tissues. Conversely, cellular models of clonal origin, such as iPSCs, enable an *in vitro* platform with minimized variability and facilitate modeling the cellular and molecular consequences of sex chromosome aneuploidies during the earliest stages of embryonic development. This also allows the systematic investigation of the impact exerted by the additional copy of Y and X-linked genes on the global transcriptome with tissue-specific resolution.

Leveraging our refined cellular model, we conducted a comprehensive transcriptional analysis encompassing JS fibroblasts, iPSCs, and NSCs juxtaposed with KS cells. This endeavor unveiled intriguing differences in the expression of most PAR1 genes, observing higher expression levels in 47,XYY compared to 47,XXY. This finding prompts a deeper investigation into whether the transcriptional contribution of the PAR gene copy on the inactive X is equivalent to the one on the Y or if it is negatively modulated by the surrounding heterochromatic environment in female and KS cells, as previously suggested^53–56^. Relevantly, our study unraveled a transcriptional feedback mechanism modulating NPX homologs in Y supernumerary cells, a phenomenon not detected in X aneuploid iPSCs. This intriguing finding underscores the need for an in-depth exploration of the epigenetic mechanisms governing such regulatory dynamics, particularly in light of our findings that the expression levels of *UTY* directly modulate *UTX*.

Furthermore, our analyses identified a shared transcriptomic signature between JS and KS, discernible already at the iPSC stage, with significant enrichment for processes related to neurological functions. This transcriptomic convergence underscores potential commonalities in the molecular pathways underpinning the pathophysiology of male sex chromosome aneuploidies. These results are particularly relevant considering that minimal DNA methylation differences exist between 47,XYY and 46,XY cells. While the well-established phenomenon of DNA methylation spreading on the supernumerary X chromosome in KS has been linked to a global transcriptional impact in primary cells^57,58^, our findings in JS iPSCs challenge this notion. Thus, the identification of highly similar patterns of differentially expressed genes in JS and KS could indicate that, in X aneuploidies, the DNA methylation of the supernumerary X could play a marginal role in regulating the global transcriptome at the pluripotent stage.

## MATERIALS AND METHODS

### Fibroblast reprogramming into iPSCs

Fibroblasts were obtained from the NIGMS Human Genetic Cell Repository at the Coriell Institute for Medical Research (**Table S1**), cultured, and expanded for at least three passages in high glucose DMEM supplemented with 15% FBS, 1% non-essential amino acids, and 1% penicillin-streptomycin before cell reprogramming. About 5 × 10^4^ –10 × 10^4^ cells/well were seeded on iMatrix-coated 6-well plates 24 h before^43^ reprogramming. The reprogramming was performed as described in. Briefly, we used the Stemgent® StemRNA-NM Reprogramming Kit to transfect the fibroblasts daily with synthetic NM-RNAs in the presence of Essential 8 Media (E8, Thermo Fisher Scientific) for four consecutive days. After ten days, emerging induced pluripotent stem cell (iPSC) colonies were manually picked and transferred into 96-well plates coated with Matrigel (Corning) in E8 with RevitaCell™ (Thermo Fisher Scientific). Individual iPSC colonies were then dissociated with Versene (Thermo Fisher Scientific), expanded, and, after ten passages, subjected to pluripotency assessment by Taqman assay and immunofluorescence.

### Human stem cell culture and transfections

The established hiPSC lines were cultured on hESC-qualified Matrigel (Corning) coated 6-well plates in E8 media and passaged with Versene in E8 supplemented with 5 μM Rock Inhibitor (Y-27632). Human iPSC lines were incubated at 37°C in 5% CO2 and 5% O2. For NPY gene silencing, 47,XYY iPSCs have been transfected with selected siRNAs (50nM; **Table S9**) using RNAiMAX Lipofectamine (Thermo Scientific). Cells were incubated in E8 media for two days after transfection before collection for RNA extraction. The hESC H1 (WA01) cell line used in this study was obtained from the WiCell biobank. hESCs were cultured on vitronectin-coated (VTN, Thermo Fisher Scientific) and E8 Medium, incubated at 37°C, 5% CO2, and passaged using Accutase (Thermo Fisher Scientific) and 5 μM Rock Inhibitor (Y-27632).

### Plasmids generation and stable transfection

The pTP6-3xFlag-KDM6A was obtained through EcoRI digestion and subcloning into the pTP6 vector^59^ of the 3xFlag-KDM6A sequence optimized for human expression (purchased from Thermo Fisher Scientific). The cDNA encoding for human ZFX was purchased from Genescript, subcloned in the pTP6-3xT7 vector through EcoRI digestion, and then amplified using the primers described in **Table S9** for subcloning inside the inducible plasmid backbone PiggyBac (System Biosciences, PB210PA-1) through NheI and XhoI double digestion.

hESCs were transfected with either pTP6-3xFlag-KDM6A or PiggyBac-3xT7-ZFX plasmids (2.5 ug) in the presence of Lipofectamine 3000 (Thermo Fischer) to generate stable H1 KDM6A and ZFX overexpressing cells. Twenty-four hours after transfection, hESCs were selected with 1mg/ml Puromycin (Thermo Fischer). Individual clones were manually picked and transferred to a 96-weel for further expansion. The inducible ZFX expression is obtained through treatment with Doxycycline at a concentration of 1ug/mL for 24 or 48 hours.

### KaryoStat analyses and DNA Fluorescence *in situ* hybridization (DNA-FISH)

A KaryoStat assay was used to accurately detect chromosomal abnormalities in the three iPSC clones obtained from the three fibroblasts’ donors. The assay, including arrays, reagents, and data analysis, was performed by Thermo Fisher Scientific.

The DNA Fluorescence in situ hybridization (DNA-FISH) was performed to validate further the Y copy number in the 47,XYY iPSC clones. iPSCs were cultured on Matrigel-coated chambered glass slides (Thermo Fisher, 177402) and fixed in a fixative solution containing 3:1 Methanol: Acetic Acid. After the Pepsin digestion at 37°C for 2-4 minutes, the slides were incubated in formaldehyde for 10 minutes and dehydrated in an ethanol series, 70%, 85%, and 100%, for 2 minutes each. The slides were then denatured in the presence of X and Y chromosome probes (CytoCell, LPE 0XYc / LPE 0XYq) on a heat block at 75C for 5 minutes and hybridized at 37°C overnight in a humidity chamber. After two washes in 1X SCC solution (Thermo Fisher, AM9763), at 37C and 45C, the slides were incubated in the presence of 4x SSC, 0.1% Tween 20 for 1 minute at room temperature, followed by an ethanol series dehydration, 70%, 85%, and 100%, for 3 minutes each. The slides were mounted with ProLong Glass antifade Mounting solution with DAPI (Thermo Fisher Scientific, P36931). DNA-FISH images were acquired as Z-stacks, processed by maximum intensity projections, and computer cleared using a DMi8 THUNDER Imager (Leica) using a 1.4 NA/100X oil immersion objective (Leica) and LAS X software.

### Differentiation of iPSCs into Neural stem cells (NSCs)

The differentiation of iPSCs into Neural stem cells was established using the PSC Neural Induction Kit from Thermo Fisher. Briefly, iPSCs were detached when they reached 70-80% confluency in small clumps using Versene and seeded at a dilution 1:6-1:8 on Matrigel-coated dishes in the presence of E8 + 10 μM Y-27632. Twenty-four hours after seeding, the Neural Induction Media (Neurobasal Medium + 1X Neural Induction Supplement) was added and used for seven days. On day 7 of neural induction, NSCs that reached P0 were harvested using Accutase, seeded at 1.0*10^6^ cells on Matrigel-coated 6-well plates, and cultured in Neural Expansion Media (50% Neurobasal Medium + 50% Advanced DMEM/F12 media + 1X Neural Induction Supplement) for about five days, until they reached 90% confluency. NSCs have been further expanded in Neural Expansion Media and collected at P2 for RNA extraction and immunostaining.

### X-Chromosome short tandem repeat (STR)

The STR analysis was performed on genomic DNA using a GlobalFiler kit (Thermo Fisher Scientific) following the manufacturer’s recommendations. A total of 24 loci were amplified, including 21 autosomal STR loci and three gender-linked loci. PCR amplicons with the internal size standard GeneScan 600 LIZ Size Standard v2.0 were loaded and resolved on SeqStudio Genetic Analyzer (Thermo Fisher Scientific). Data were analyzed with GeneMapper ID-X software v1.6 (Applied Biosystems).

### Immunofluorescence and image acquisitions

iPSCs were plated on Matrigel-coated coverslips and fixed for 12 min with 3% paraformaldehyde for immunostaining 72 hours after seeding. Fixed cells were permeabilized with 0.25% Triton-X100 in PBS, incubated overnight with primary antibodies for pluripotency markers (**Table S8**), washed, incubated with secondary antibodies, and mounted with ProLong Glass antifade Mounting solution with DAPI (Thermo Fisher Scientific, P36931). Immunostainings for pluripotency markers were acquired using an EVOS^TM^ FL Auto 2 Imaging System (Thermo Fisher Scientific) using a 1.30 NA/40X oil immersion objective (Olympus).

### RNA extraction and qPCR

The total RNA extraction from iPSCs, fibroblasts, and NSCs was performed using the KingFisher Duo Prime benchtop sample extraction instrument (Thermo Fisher Scientific) using MagMAX mirVana Total RNA Isolation Kit (Applied Biosystems) following the manufacturer’s instructions. The cDNA was synthesized with the SuperScript VILO IV cDNA Synthesis Kit (Thermo Fisher Scientific). Gene expression was determined by real-time qPCR on a QuantStudio 3 Real-Time PCR System (Thermo Fisher Scientific) using TaqMan™ Fast Advanced Master Mix (Thermo Fisher Scientific) and TaqMan® Gene Expression Probes (**Table S9**). Individual gene expression was normalized on Tata Binding Protein (TBP) using the 2-ÄÄCt 2-(Delta Delta C(T)) as a relative quantification method.

### Bulk RNA-Seq library preparation and sequencing

RNA libraries were generated using the human mRNA TruSeq Stranded library preparation KIT from Illumina and profiled using NovaSeq6000 and NovaSeq XPlus systems with 150 bp paired-end sequencing method. An average of 30M reads were obtained for each sample. Samples with less than 16M input reads and lower than 75% assigned reads were removed from the analysis (**Table S10 and Table S11**).

### RNA-seq data processing and analysis

To assess the quality of the raw RNA-seq data, we used the FastQC tool (http://www.bioinformatics.babraham.ac.uk/projects/fastqc/). This was followed by trimming adapter sequences, filtering out low-quality bases with a minimum quality score of 25, and discarding reads shorter than 35 bases with BBDuk (BBmap suite v37.62). Then, we performed a quality reassessment of the trimming step with FastQC. The preprocessed high-quality reads were then aligned to the GRCh38.101 release of the Ensembl human reference genome using the STAR software v. 2.7.10^60^. To maintain high precision in the alignment process, we permitted a maximum of three mismatches per read (--outFilterMismatchNmax 3) and limited the mismatch ratio to no more than 10% of the read length (--outFilterMismatchNoverLmax 0.1). The alignment quality was further ensured by requiring that the alignment score be at least 66% of the maximum possible score based on the read length (--outFilterScoreMinOverLread 0.66), effectively filtering out lower-quality alignments. We used featureCounts (Subread suite v. 2.0.2)^61^ to quantify gene expression levels. To address potential batch effects, we used ComBat-seq^62^. We divided the adjusted count matrix per cell type to independently perform the gene filtering and differential expression analysis. Genes with low expression were removed from the analysis using the HTS Filter package^63^. Differential gene expression analysis was conducted via the DESeq2 package^64^ in R. We applied a false discovery rate (FDR) cutoff of less than 0.05 and a |Log_2_FC| > 0.58 threshold to consider a gene differentially expressed. An FDR<0.01 threshold was applied for specific analysis and plots when specified in the figure legends.

### Identification of genes with dosage sensitivity to chromosome Y copy number

We used the count matrix for the iPSCs samples to perform the correlation analysis. We applied batch correction with ComBat-seq, removed low-expressed genes with the HTS Filter package, calculated normalized values using DESeq2, and then performed a Pearson correlation for every expressed gene (Y ∼ E, where Y is the number of Y chromosomes and E is the normalized expression).

### Around the X Chromosome

Healthy male (XY) samples (n=34) were used as a reference to calculate the median FPKM level of each gene (male median) after removing non-expressed genes as previously described^65^. Additionally, n=6 female, n=24 KS, and n=37 JS iPSC samples were plotted. Next, for every sample, the FPKM level of each gene was divided by the male median. These fold changes were then plotted in a moving average plot using a loess fit, adjusting the span to 0.45, and determining the sliding window size as a proportion of observations in each local regression.

MM (Male median) = Median (FPKM in reference XY samples) of a specific gene FC (Fold Change) = FPKM in each sample/MM of a specific gene

### Whole Exome sequencing and Allele-Specific Expression analysis

According to manufacturer instructions, genomic DNA was extracted from hiPSC lines using the DNeasy Blood & Tissue Kits (Qiagen). The DNA quality was assessed using the TapeStation Agilent Genomic DNA ScreenTape Assay (Agilent) before DNA shipment. Beijing Novogene Bioinformatics Technology, Co., Ltd. (Beijing, China) performed the WES analysis. Target enrichment was performed to construct the exome library using the Agilent SureSelect Human All Exon V6 kit (Agilent Technologies, Inc., Santa Clara, CA, United States), according to the manufacturer’s protocol, and sequenced on Illumina Platform. An effective coverage of around 100× was obtained for all samples. Paired-end sequence reads for each individual were aligned to the GRCh38 reference human genome sequence using the sentieon bwa mem (BWA v0.7.17-r1188 aligner). Then, aligned reads were sorted, duplicates were marked using sentieon-Dedup, sorted with sentieon-sort, and base quality recalibrated using sention-ReadWriter (sention-genomics v202112.02), respectively. For the Variant Calling, a combination of GATK v4.2.6.1 SelectVariants, followed by sentieon TNfilter. We utilized the Sentineons’ TNhaplotyper2 algorithm with specific options to enhance accuracy.

The --call_germline_sites option directed the algorithm to call germline variants, while the --call_pon_sites option ensured using a panel of normal samples from the 1000 Genomes Project to classify common variants. Finally, bcftools 1.11+htslib-1.11 was used to keep the variants that met the following criteria: the FILTER field set as PASS or panel of normals and depth of coverage > 10. To ensure the accuracy of the called variants, we applied an additional filter to remove variants with no reads supporting either the reference or alternate alleles. We used ASERead Counter from GATK v4.2.2.0 for ASE analysis to obtain read counts covering each allele on SNPs. To detect SNPs differing from the reference genome, we set stringent parameters of base quality BQ ≥ 30, mapping quality MQ ≥ 10, and read depth > 6. We excluded intronic sites and non-uniquely mapped genes. Reference allele expression was calculated as reference allele counts divided by total counts. SNPs were classified as biallelic if reference allele expression was between 0.1 and 0.9 and monoallelic if outside this range. Genes were considered biallelic if at least one SNP in the gene was biallelic. Importantly, we detected a single biallelic expressed SNP on the X chromosome in sample JS2 #A corresponding to the gene NUDT10. We took 50 bases upstream and downstream of the SNP and performed a blast against the RefSeq RNA database to investigate how this SNP could be found biallelic in a sample with only one X. We also visualized the bigWig files using the genomic viewer IGV^66^. We found a 100% match of the sequence against NUDT10 and a 99% with one mismatch against NUDT11. We concluded that the identified SNP, attributable to NUDT11, was mistakenly mapped to NUDT10, possibly due to a poorly mapping artifact resulting in a false SNP in one JS2 iPSC clone.

### Reduced-representation bisulfite (RRBS) library preparation and sequencing

The gDNA was extracted using the DNeasy blood and tissue genomic DNA extraction kit from Qiagen®. Beijing Novogene Bioinformatics Technology, Co., Ltd (Beijing, China) generated the DNA libraries. Then, the gDNA was sequenced using Illumina Novaseq6000 or NovaseqX 2 x 150 bp.

### RRBS data analysis

The quality of raw data was assessed using FastQC. We then proceeded with adapter trimming C using trimGalore v.0.6.6 to prepare the DNA methylation data. After a quality reassessment using FastQC, the high-quality reads were aligned to the GRCh38.105 release of the Ensembl human reference genome using Bismark v.0.23.1. CpG site methylation extraction was conducted with MethylDackel v 0.6.1. Then, differential methylation analysis was performed with the DSS package in R^67^. We defined differentially methylated regions (DMRs) by a minimum length of 50 base pairs and a minimum of three CpGs. DMRs in a distance within 50 base pairs of one another were merged. We established a significance threshold for loci at 1e-5, requiring that DMRs contain at least 50% differentially methylated CpGs throughout their entire length after the merging.

## DATA AVAILABILITY

The RNA-seq and RRBS-seq data generated in this study are available in the Gene Expression Omnibus (GEO) database under accession numbers GSE264395 and GSE264396, respectively. The RNA-seq data from our previous studies that we have reanalyzed in this study are publicly available at GEO under accession numbers GSE152001 and GSE220268.

## FIGURE TITLES AND LEGENDS

**Figure S1: 47,XYY Fibroblasts reprogramming into iPSCs. A)** mRNA expression of the *UTY*, *ZFY,* and *KDM5D* Y-linked genes by Taqman assay in 47,XYY and 46,XY fibroblasts. Values are normalized on the housekeeping gene TATA box-binding protein (*TBP*). Bars are the average ± std of three independent experiments for each iPSC clone. Values from two 46,XY control’s fibroblasts were merged (n=6) to calculate the statistic using a Student’s t-test, one-tailed distribution, two-sample equal variance. HC, healthy controls 46,XY. **B)** Schematic of the mRNA non-integrative reprogramming method used in the study.

**Figure S2: Validation by DNA-FISH of Y chromosome copy gain in iPSCs.** Top: Representative DNA-FISH images showing the X chromosome (green) and Y chromosome (red) in JS-iPSCs. DNA is stained with DAPI (blue). Scale bar = 50 µm. Bottom: Magnified view of the dashed square in the top image. Scale bar = 5 µm.

**Figure S3: Karyostat arrays on 47-XXY iPSCs. A)** Summary of the validated iPSCs. The probe binding to Xp22.33 also recognizes the Yp11.2 region and corresponds to the PAR1 territory on X and Y chromosome, respectively; therefore, the showed X chromosome’s duplication for this region is an artifact of the probe binding to the PAR1 region of both sex chromosomes. Thus, the results show two copy number state (CN = 2) for the whole chromosome Y except for the Yp11.2 probe (CN = 1) automatically assigned to the X chromosome (Xp22.33, CN = 3). **B)** KaryoStat+ whole genome view. The pink, green, and yellow colors indicate the raw signal for each chromosome probe, while the blue represents the normalized probe signal, which is used to identify copy numbers and aberrations. Black arrows indicate chromosome gains.

**Figure S4: Pluripotency characterization by immunostaining on 47,XYY iPSCs generated in this study. A)** Transcript levels of the pluripotency markers *SOX2*, *NANOG,* and *OCT4* in the iPSCs generated in the study analyzed by TaqMan Assay. Bars are the average ± std of three independent experiments. Values from two 46,XY control’s iPSCs were merged (n=6) to calculate the statistic using a Student’s t-test, one-tailed distribution, and two-sample equal variance. *p<0.05, **p<0.01, n.s. = not significant. **B-C)** Staining for the pluripotency markers NANOG (red), SOX2 (red), and SSEA4 (green). See **Table S8** for antibody information. DAPI (blue) is used as a counterstain to detect nuclei. Images have been acquired using an EVOS^TM^ FL Auto 2 Imaging System (Thermo Fisher Scientific) equipped with a 1.30NA/40X oil immersion objective (Olympus). Scale bars, 50 μm.

**Figure S5: Derivation of neural stem cells from 47,XYY and 46,XY iPSCs. A)** Timeline of iPSCs differentiation into NSCs. At approximately 20-22 days of differentiation, NSCs reached passage 2 (P2) and were collected for RNA profiling or Immunofluorescence. **B-C)** Taqman assay showing the mRNA expression of the pluripotency marker *OCT4*, and NSC markers *NESTIN* (*NES*) and *TUBULIN3* (*TUBB3*) in 46,XY (B) or 47,XYY cells (C). Values are normalized to the internal control TATA-binding protein (*TBP*). Each Purple dot represents an independent qPCR sample. One-way ANOVA and post-hoc Tukey HSD analyses were used to test the significance between groups. P values are shown. **D)** 47,XYY NSCs stained for the indicated markers. Images have been acquired using an EVOS^TM^ FL Auto 2 Imaging System equipped with a 0.3NA/10X oil immersion objective (Olympus). Scale bars, 200 μm. See

**Table S8** for antibody information.

**Figure S6: Allele-specific expression (ASE) analysis shows the biallelic expression of SNPs at the PAR region in 47,XYY cells**. Scatter plot profiles of coupled WES analysis and allele-specific RNA-Seq analysis performed on autosomal (upper panel) and X chromosomes (lower panel) showing the mono-(orange dots) or biallelic (light blue dots) gene expression status in 47,XYY iPSCs. Gray rectangles indicate PAR1 and PAR2 regions, respectively. Solid dots indicate non-PAR genes; open dots show PAR genes.

**Figure S7: Signature of cell-type specific PAR expression.** Box Plot of normalized TMM expression of PAR genes in control 46,XY and 47,XYY Fibroblasts, iPSCs, and NSCs. Each purple dot represents an independent RNA sample. Two-way ANOVA and post-hoc Tukey HSD analyses were used to test the significance of the groups and karyotypes. *p_adj_ < 0.05; ** p_adj_ < 0.01; *** p_adj_ < 0.001.

**Figure S8: Genome-wide effects of chromosome Y aneuploidy. A-C)** Volcano plots effect size and significance of DEGs detected in 47,XYY versus 46,XY (A) Fibroblasts, (B) iPSCs, and (C) NSCs. The numbers of up and down-regulated DEGs are shown for each contrast. **D)** Summary of the autosomal, PAR, and non-PAR X and Y DEGs detected in the three cell types. FDR<0.05 and Log_2_FC >|0.58|.

**Figure S9: Overdosage of PAR and NPY genes in Chromosome Y aneuploid cells. A-C)** Heatmap showing the TMM expression of upregulated NPY DEGs in 47,XYY vs. 46,XY contrast in (A) Fibroblasts, (B) iPSCs, and (C) NSCs. **D-F)** Heatmap showing the TMM expression of upregulated PAR DEGs in (D) Fibroblasts, (E) iPSCs, and (F) NSCs.

**Figure S10: Y-linked gene overdosage modulates X chromosome expression. A-B)** Heatmap showing the TMM expression of up- and downregulated NPX DEGs in 47,XYY (A) Fibroblasts, (B) NSCs, and (C) iPSCs. **D-F)** Venn diagrams of shared NPX DEGs in (D) fibroblasts and iPSCs, (E) iPSCs and NSCs, or (F) the three cell types. The number of NPX genes is shown in each diagram.

**Figure S11: Methylation analysis in 47,XYY A)** PCA of methylation sites in fibroblasts and iPSCs on the whole genome and **B)** Number, location, and direction of DMRs in fibroblasts and iPSCs.

**Figure S12: Box plot of the TMM Normalized PAR gene expression in control 46,XY, 47,XYY, and 47,XXY Fibroblasts.** Each Purple dot represents an independent RNA sample. One-way ANOVA and post-hoc Tukey HSD analyses were used to test the significance between karyotypes. *p_adj_ < 0.05; ** p_adj_ < 0.01; *** p_adj_ < 0.001.

**Table S1**: Description of Jacob syndrome patient’ cohort

**Table S2**: List of informative STRs unequivocally assigned to each fibroblast patient and corresponding iPSC clones.

**Table S3**: List of expressed genes in 47,XYY vs. 46,XY Fs, iPSCs, and NSCs.

**Table S4**: List of DEGs in 47,XYY vs. 46,XY Fs, iPSCs, and NSCs with FDR< 0.05 and FDR<0.05 and Log_2_FC >|0.58| and FDR <0.01.

**Table S5**: Pearson correlation analysis in iPSCs. List of genes sensitive to Y chromosome number.

**Table S6**: List of DMRs identified in the contrast 47,XYY vs 46,XY fibroblasts and iPSCs.

**Table S7**: List of shared DEGs in the contrasts 47,XYY vs. 46,XY and 47,XXY vs 46,XY in Fibroblasts and iPSCs. FDR<0.05 and Log_2_FC >|0.58|.

**Table S8**: List of antibodies used in the study.

**Table S9**: List of Taqman probes, oligos, and siRNAs used in the study

**Table S10:** List of RNA-seq libraries used in this study and NCBI GEO accession numbers.

**Table S11:** List of RNA-seq libraries sequenced and mapping quality.

## AUTHOR CONTRIBUTIONS

VA performed somatic cell reprogramming. VA and LCM cultured the hiPSC lines, differentiated derivatives, and prepared the genomic DNA (gDNA) and RNAs used in this study. RA performed STR genotyping, gDNA, RNA extraction, and RNA-SEQ library preparation. KC-L, JD-C, and SR performed transcriptomic and methylome analyses. GR performed DNA-FISH and immunofluorescence staining, as well as Karyostat sample preparation. FK performed the experiments on ZFX and UTX overexpressing cell lines. VA and AA wrote the manuscript. VA, KC-L, LCM, RA, GR, FK, JD-C, SR, and AA revised and edited the manuscript. AA conceived, designed, and supervised the study.

## ACKNOWLEDGMENTS

We thank the genomic unit of the KAUST BioCoreLab for technical support with RNA-Seq libraries’ quality control. We thank the KAUST imaging and characterization facility for support with image acquisition. We thank Maryam Alowaysi and Juan Pablo Padilla Martinez for technical support with the pTP6-3xFlag-KDM6A overexpressing cell line. This work was funded by baseline funding (BAS 1077-01-01) to AA.

## COMPETING FINANCIAL INTERESTS

The authors declare no competing financial interests

## Notes

### Competing Interest Statement

The authors have declared no competing interest.

