## Supplemental_Figures for "An iPSC-based model of Jacob Syndrome reveals a DNA methylation-independent transcriptional dysregulation shared with X aneuploid cells"

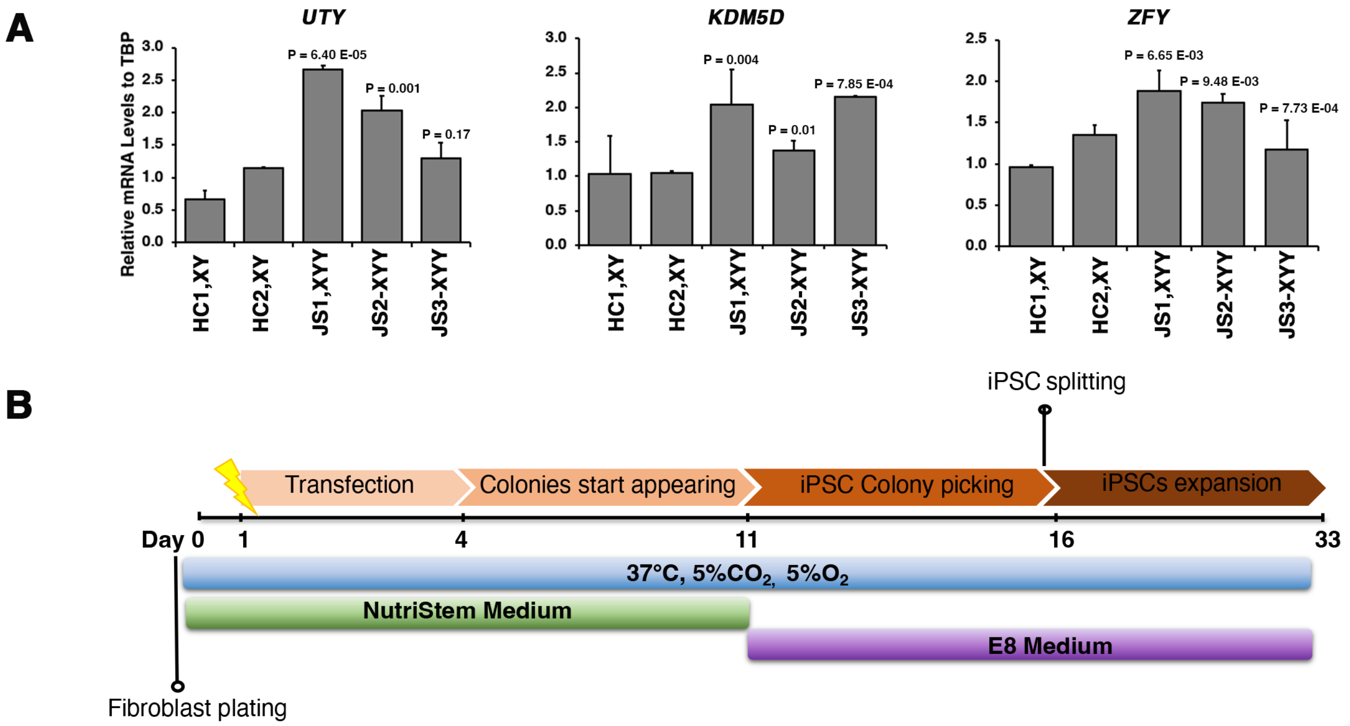
A
UTY
KDM5D
ZFY
Relative mRNA Levels to TBP
P = 6.40 E-05
P = 0.001
P = 0.17
P = 0.004
P = 0.01
P = 7.85 E-04
P = 6.65 E-03
P = 9.48 E-03
P = 7.73 E-04
HC1,XY
HC2,XY
JS1,XYY
JS2-XYY
JS3-XYY
B
iPSC splitting
Transfection
Colonies start appearing
iPSC Colony picking
iPSCs expansion
Day 0
1
4
11
16
33
37°C, 5%CO2, 5%O2
NutriStem Medium
E8 Medium
Fibroblast plating

### Figure S1

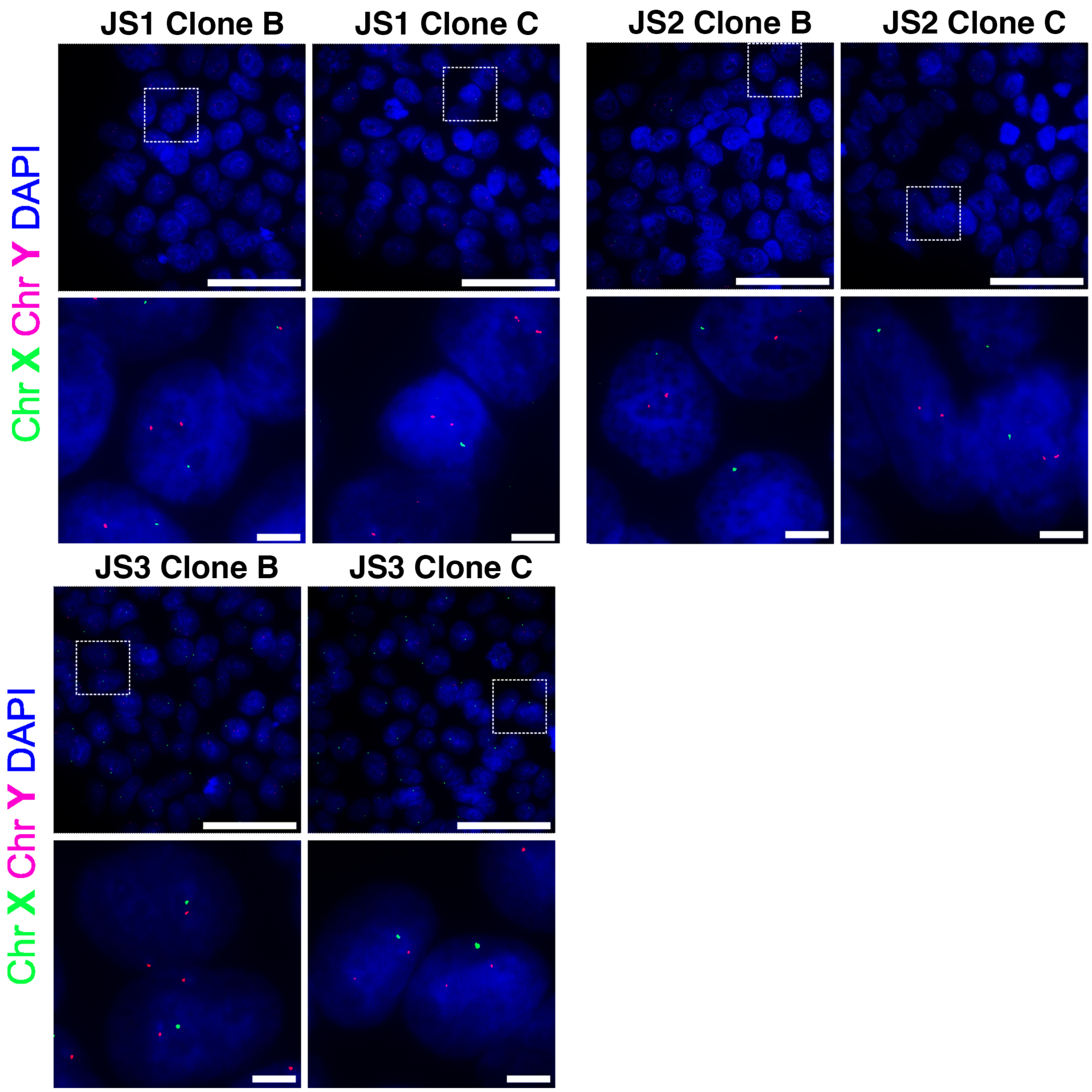
JS1 Clone B
JS1 Clone C
JS2 Clone B
JS2 Clone C
Chr X Chr Y DAPI
JS3 Clone B
JS3 Clone C
Chr X Chr Y DAPI

**Figure S2**

**A**

| Clone name | Chromosome | Type | Cytoband Start | CN State | Size (kbp) |
|---|---|---|---|---|---|
| JS1 Clone A | Y | Gain | p11.2 | 2.00 | 18,642 |
| | Y-X PAR1 | Gain | p22.33 | 3.00 | 2,111 |
| JS1 Clone B | Y | Gain | p11.2 | 2.00 | 19,068 |
| | Y-X PAR1 | Gain | p22.33 | 3.00 | 2,035 |
| | 11 | Gain | q13.5 | 3.00 | 1,218 |
| JS1 Clone C | Y | Gain | p11.2 | 2.00 | 18,604 |
| | Y-X PAR1 | Gain | p22.33 | 3.00 | 1,504 |
| JS2 Clone A | Y | Gain | p11.2 | 2.00 | 23,624 |
| | Y-X PAR1 | Gain | p22.33 | 3.00 | 2,111 |
| JS2 Clone B | Y | Gain | p11.2 | 2.00 | 23,646 |
| | Y-X PAR1 | Gain | p22.33 | 3.00 | 2,525 |
| | 6 | Gain | q25.1 | 3.00 | 1,998 |
| JS2 Clone C | Y | Gain | p11.2 | 2.00 | 23,615 |
| | Y-X PAR1 | Gain | p22.33 | 3.00 | 1,513 |
| JS3 Clone A | Y | Gain | p11.2 | 2.00 | 21,481 |
| | Y-X PAR1 | Gain | p22.33 | 3.00 | 1,643 |
| JS3 Clone B | Y | Gain | p11.2 | 2.00 | 19,068 |
| | Y-X PAR1 | Gain | p22.33 | 3.00 | 1,818 |
| JS3 Clone C | Y | Gain | p11.2 | 2.00 | 21,481 |
| | Y-X PAR1 | Gain | p22.33 | 3.00 | 1,643 |

**B**

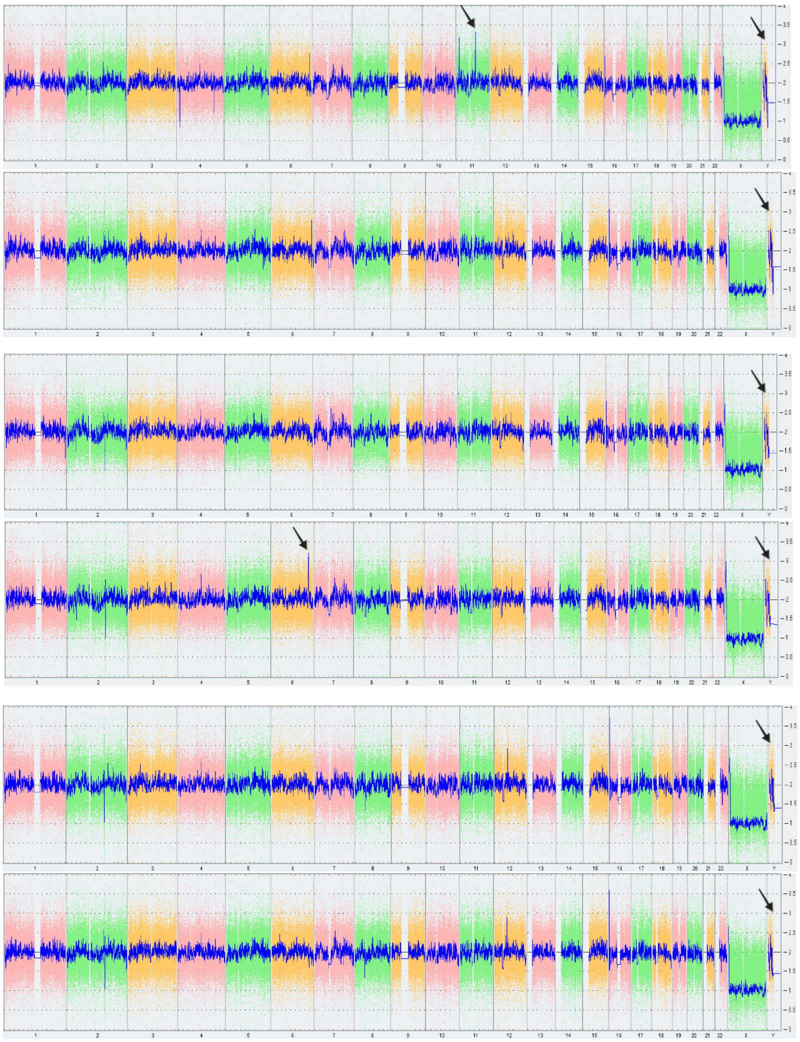

**iPSCs JS1 Clone B**

**iPSCs JS1 Clone C**

**iPSCs JS2 Clone B**

**iPSCs JS2 Clone C**

**iPSCs JS3 Clone B**

**iPSCs JS3 Clone C**

**Figure S3**

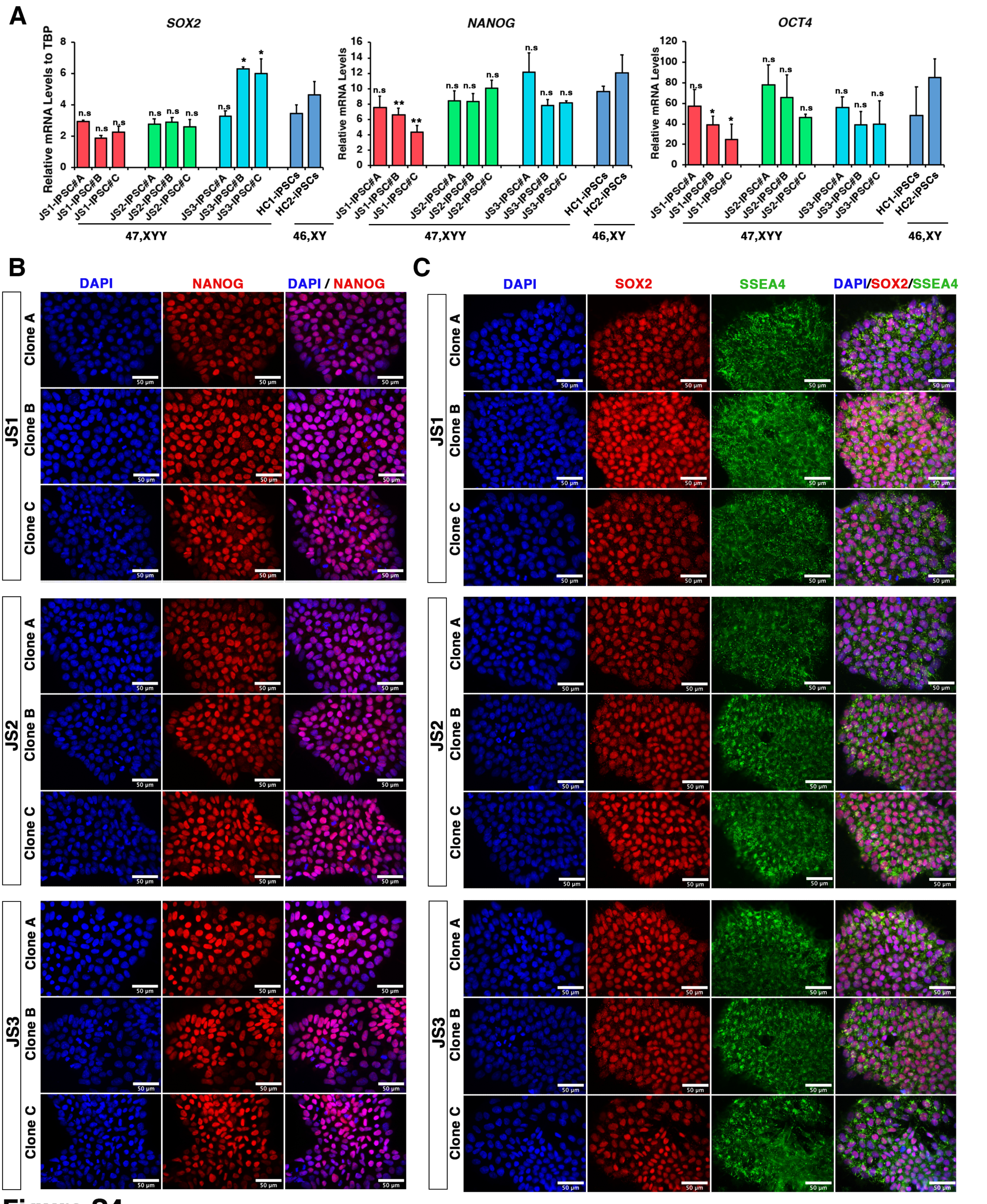
A
SOX2
NANOG
OCT4
Relative mRNA Levels to TBP
Relative mRNA Levels
JS1-IPSC#A
JS1-IPSC#B
JS1-IPSC#C
JS2-IPSC#A
JS2-IPSC#B
JS2-IPSC#C
JS3-IPSC#A
JS3-IPSC#B
JS3-IPSC#C
HC1-IPSCs
HC2-IPSCs
47,XYY
46,XY
B
DAPI
NANOG
DAPI / NANOG
C
SOX2
SSEA4
DAPI/SOX2/SSEA4
JS1
JS2
JS3
Clone A
Clone B
Clone C
50 µm

**Figure S4**

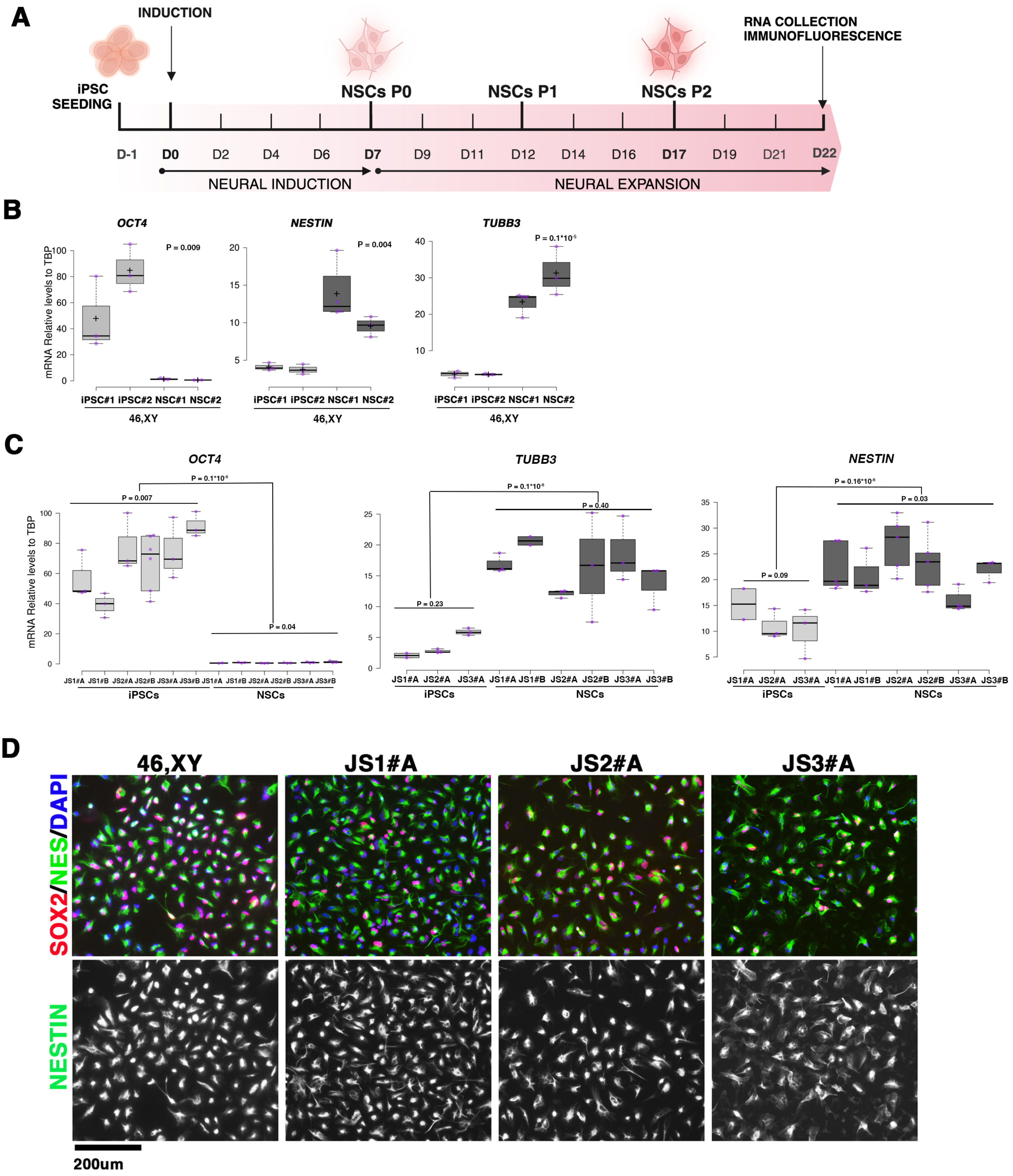
A
INDUCTION
RNA COLLECTION
IMMUNOFLUORESCENCE
iPSC
SEEDING
NSCs P0
NSCs P1
NSCs P2
D-1 D0 D2 D4 D6 D7 D9 D11 D12 D14 D16 D17 D19 D21 D22
NEURAL INDUCTION
NEURAL EXPANSION
B
OCT4
NESTIN
TUBB3
mRNA Relative levels to TBP
P = 0.009
P = 0.004
P = 0.1*10-5
iPSC#1 iPSC#2 NSC#1 NSC#2
46,XY
C
OCT4
TUBB3
NESTIN
P = 0.1*10-5
P = 0.007
P = 0.04
P = 0.23
P = 0.40
P = 0.16*10-5
P = 0.03
P = 0.09
JS1#A JS1#B JS2#A JS2#B JS3#A JS3#B
iPSCs
NSCs
D
46,XY
JS1#A
JS2#A
JS3#A
SOX2/NES/DAPI
NESTIN
200um

### Figure S5

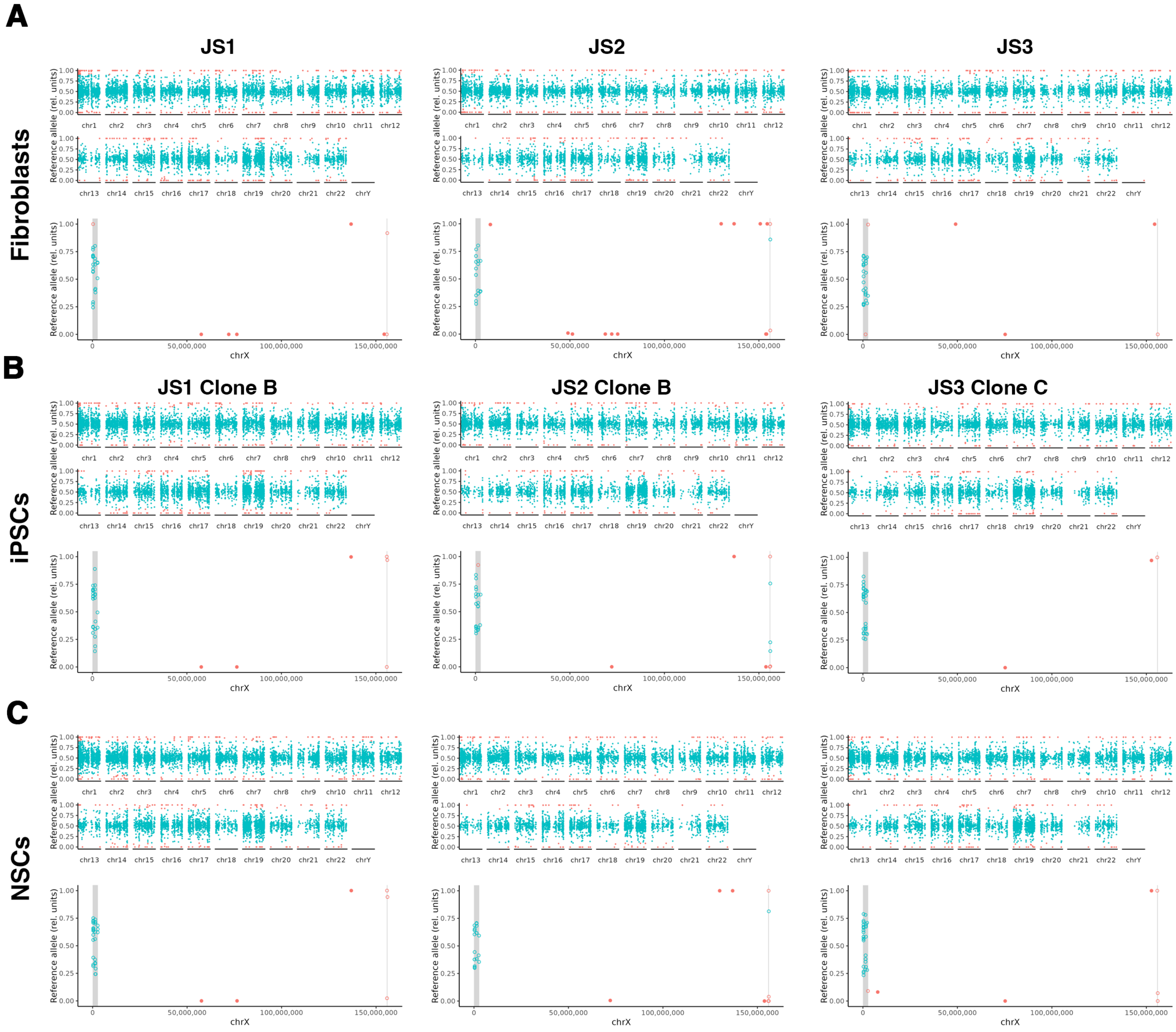
A
JS1
JS2
JS3
Fibroblasts
B
JS1 Clone B
JS2 Clone B
JS3 Clone C
iPSCs
C
NSCs
Reference allele (rel. units)
chr1 chr2 chr3 chr4 chr5 chr6 chr7 chr8 chr9 chr10 chr11 chr12
chr13 chr14 chr15 chr16 chr17 chr18 chr19 chr20 chr21 chr22 chrY
chrX
0 50,000,000 100,000,000 150,000,000
0.00 0.25 0.50 0.75 1.00

### Figure S6

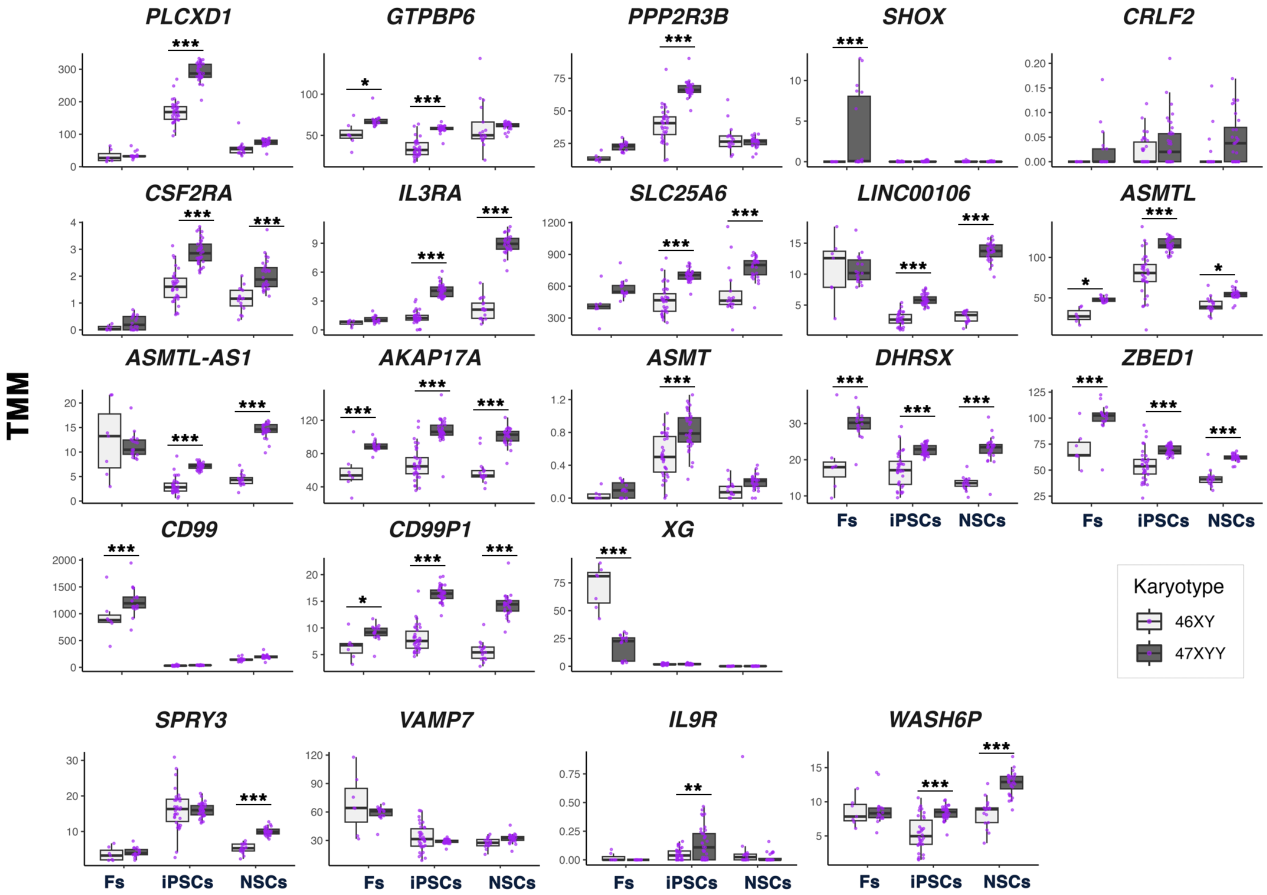

PLCXD1
GTPBP6
PPP2R3B
SHOX
CRLF2
CSF2RA
IL3RA
SLC25A6
LINC00106
ASMTL
ASMTL-AS1
AKAP17A
ASMT
DHRSX
ZBED1
CD99
CD99P1
XG
SPRY3
VAMP7
IL9R
WASH6P
TMM
Fs
iPSCs
NSCs
Karyotype
46XY
47XYY


### Figure S7

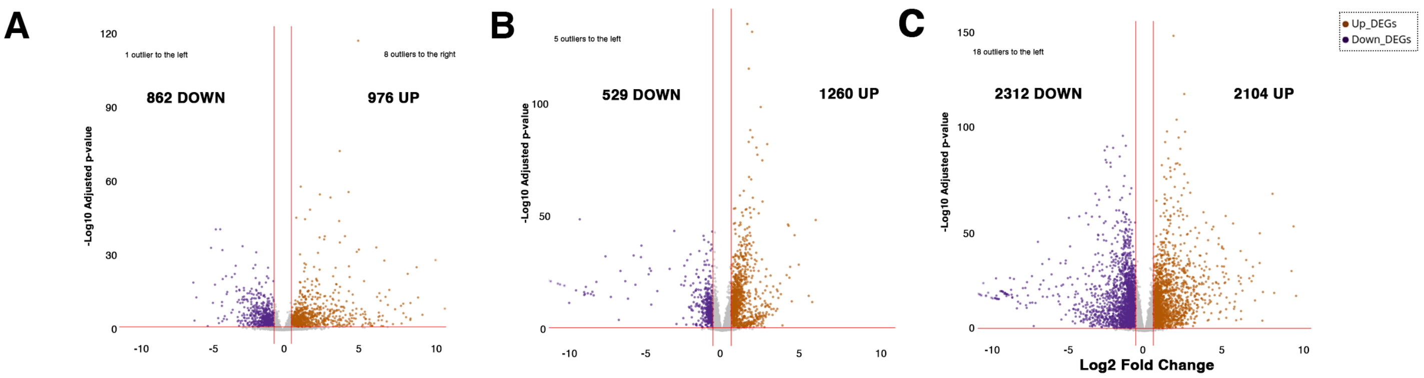


**D**

| | 47,XYY Vs. 46,XY (N.DEGs) | | |
|---|---|---|---|
| | **Fibroblasts** | **iPSCs** | **NSCs** |
| **Autosomes** | 1769 | 1696 | 4229 |
| **Sex Chromosomes** | 69 | 89 | 181 |
| **X Chromosome (non-PAR)** | 50 | 62 | 162 |
| **Y chromosome (non-PAR)** | 14 | 15 | 12 |
| **X and Y PAR genes** | 5 | 12 | 7 |

### Figure S8

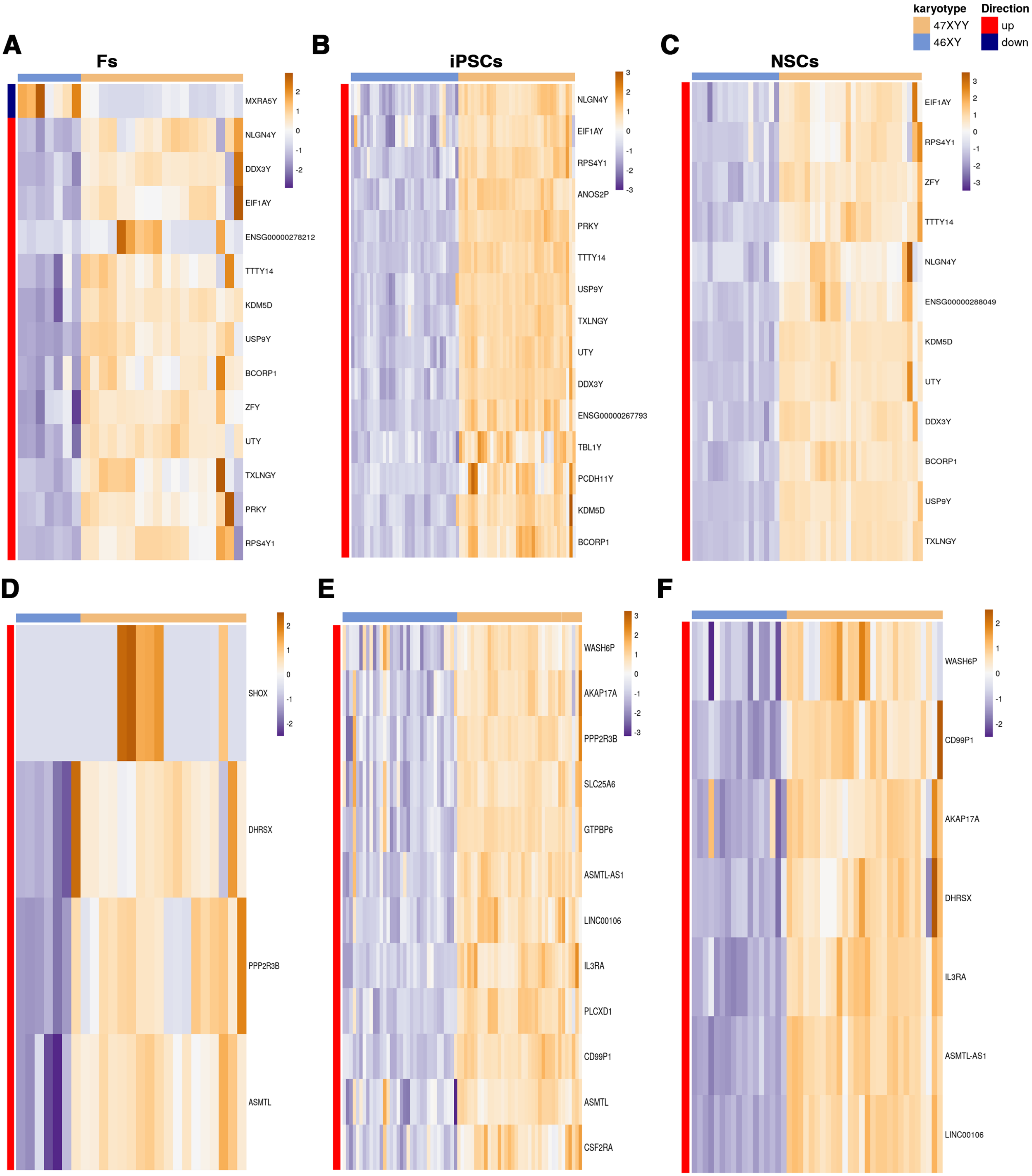
karyotype
47XXY
46XY
Direction
up
down
A
Fs
MXRA5Y
NLGN4Y
DDX3Y
EIF1AY
ENSG00000278212
TTTY14
KDM5D
USP9Y
BCORP1
ZFY
UTY
TXLNGY
PRKY
RPS4Y1
B
iPSCs
NLGN4Y
EIF1AY
RPS4Y1
ANOS2P
PRKY
TTTY14
USP9Y
TXLNGY
UTY
DDX3Y
ENSG00000267793
TBL1Y
PCDH11Y
KDM5D
BCORP1
C
NSCs
EIF1AY
RPS4Y1
ZFY
TTTY14
NLGN4Y
ENSG00000288049
KDM5D
UTY
DDX3Y
BCORP1
USP9Y
TXLNGY
D
SHOX
DHRSX
PPP2R3B
ASMTL
E
WASH6P
AKAP17A
PPP2R3B
SLC25A6
GTPBP6
ASMTL-AS1
LINC00106
IL3RA
PLCXD1
CD99P1
ASMTL
CSF2RA
F
WASH6P
CD99P1
AKAP17A
DHRSX
IL3RA
ASMTL-AS1
LINC00106

**Figure S9**

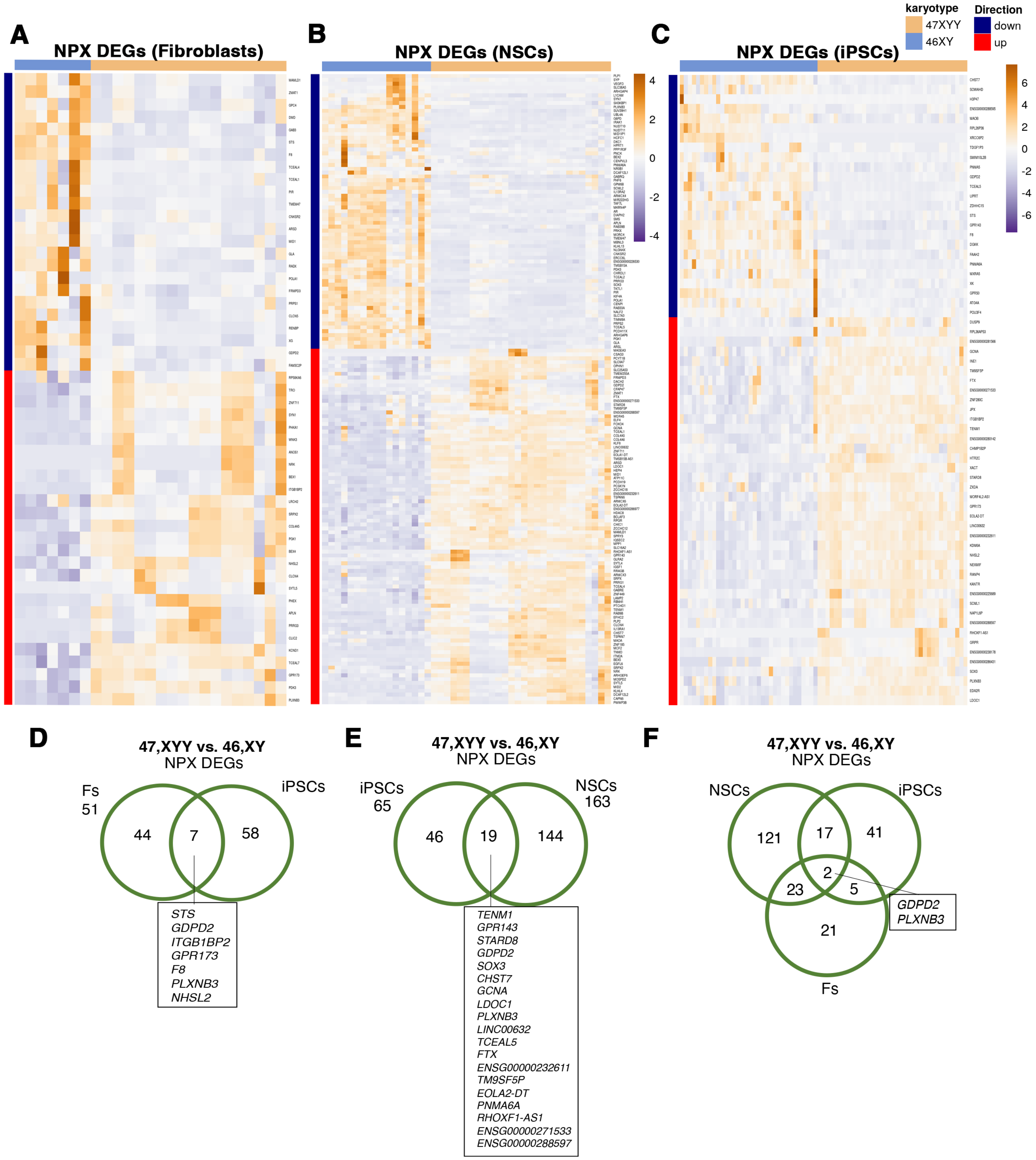

A
NPX DEGs (Fibroblasts)
B
NPX DEGs (NSCs)
C
NPX DEGs (iPSCs)
karyotype
47XXY
46XY
Direction
down
up
D
47,XYY vs. 46,XY
NPX DEGs
Fs
51
iPSCs
44
7
58
STS
GDPD2
ITGB1BP2
GPR173
F8
PLXNB3
NHSL2
E
47,XYY vs. 46,XY
NPX DEGs
iPSCs
65
NSCs
163
46
19
144
TENM1
GPR143
STARD8
GDPD2
SOX3
CHST7
GCNA
LDOC1
PLXNB3
LINC00632
TCEAL5
FTX
ENSG00000232611
TM9SF5P
EOLA2-DT
PNMA6A
RHOXF1-AS1
ENSG00000271533
ENSG00000288597
F
47,XYY vs. 46,XY
NPX DEGs
NSCs
iPSCs
121
17
41
2
23
5
21
Fs
GDPD2
PLXNB3


**Figure S10**

A

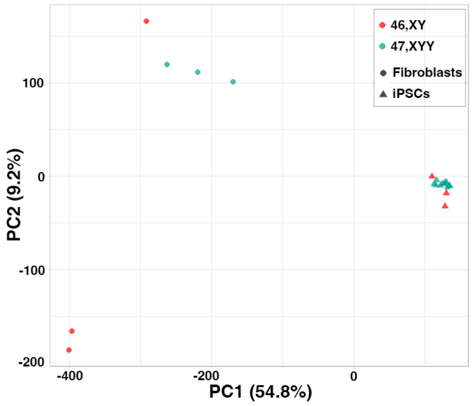


B

| Cell type | Gene Location | Hyper_total | Hypo_total |
|---|---|---|---|
| Fibroblasts | Autosome | 3384 | 691 |
| Fibroblasts | X Chromosome | 93 | 21 |
| Fibroblasts | Y Chromosome | 8 | 0 |
| iPSCs | Autosome | 28 | 31 |
| iPSCs | X Chromosome | 2 | 0 |
| iPSCs | Y Chromosome | 0 | 2 |

### Figure S11

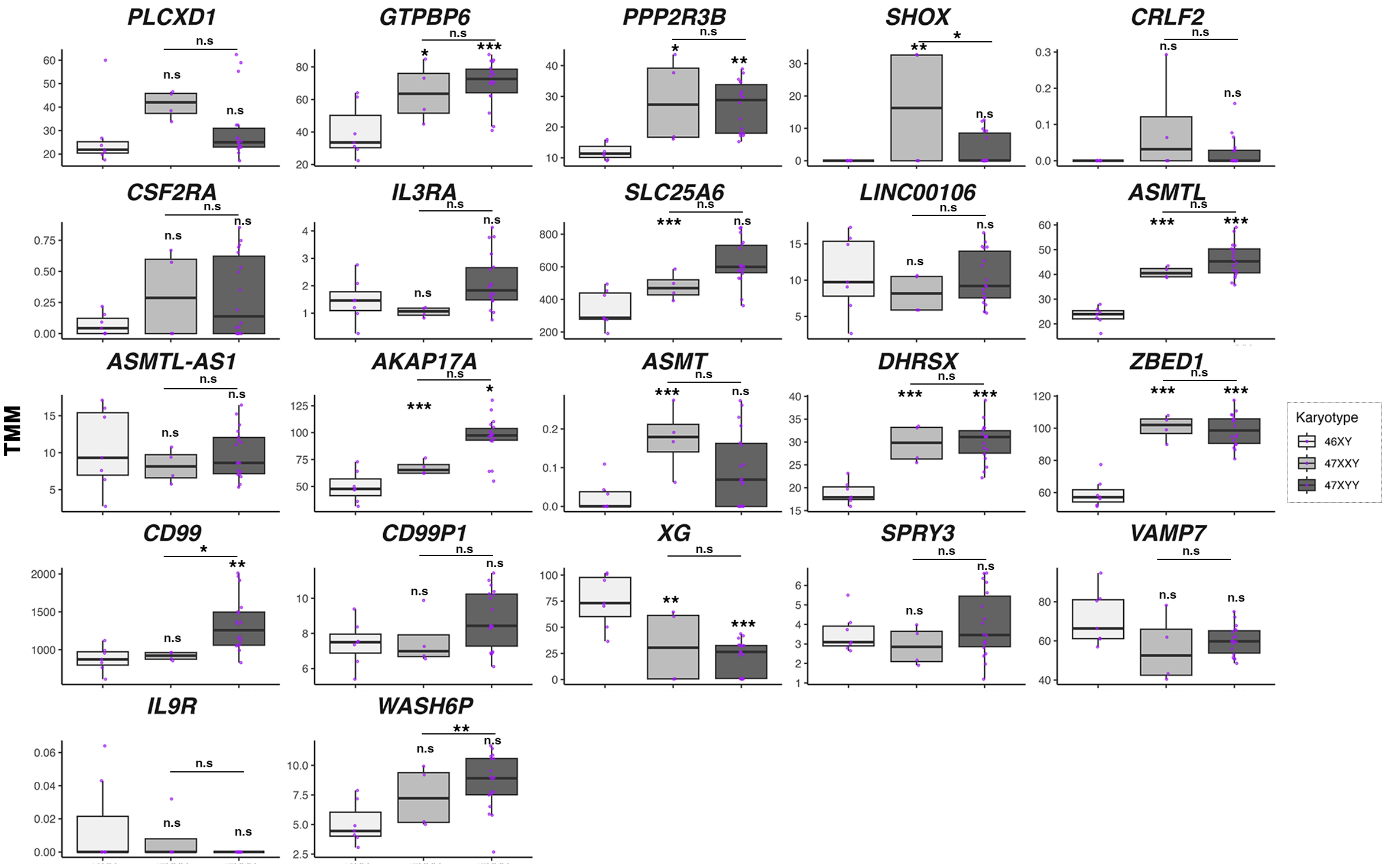
PLCXD1
GTPBP6
PPP2R3B
SHOX
CRLF2
CSF2RA
IL3RA
SLC25A6
LINC00106
ASMTL
ASMTL-AS1
AKAP17A
ASMT
DHRSX
ZBED1
CD99
CD99P1
XG
SPRY3
VAMP7
IL9R
WASH6P
TMM
Karyotype
46XY
47XXY
47XYY

### Figure S12
