## Supplementary material for "An iPSC-based model of Jacob Syndrome reveals a DNA methylation-independent transcriptional dysregulation shared with X aneuploid cells": Table S1

| **Fibroblast cell line** | **Patient age (at sampling)** | **Ethnicity** | **Remarks** | **Repository** | **Name in the paper** | **Karyotype** |
| --- | --- | --- | --- | --- | --- | --- |
| GM01250 | 23 YR | Black/African American | Phenotypically normal | NIGMS Human Genetic | JS1 | 47,XYY |
| GM09326 | 2 Months | Irish/English | Phenotypically normal | NIGMS Human Genetic | JS2 | 47,XYY |
| GM11337 | 19 FW | N/A | Induced abortion | NIGMS Human Genetic | JS3 | 47,XYY |

**Table S1**: Fibroblast lines reprogrammed in this study.
