## Supplementary material for "An iPSC-based model of Jacob Syndrome reveals a DNA methylation-independent transcriptional dysregulation shared with X aneuploid cells": Table S8

**Antibodies used for Immunofluorescence and Western Blot.**

| **Antibody** | **Species** | **Dilution** | **Manufacturer** |
| --- | --- | --- | --- |
| NANOG | Rabbit | 1:200 | Abcam Cat# ab109250 |
| OCT4 | Mouse monoclonal | 1:100 | Thermo Fisher Scientific, Cat#MA1-104 |
| SOX2 | Mouse anti Rabbit | 1:500 | Cell Signalling Technology  Cat#46743S |
| SSEA4 | Mouse monoclonal | 1:100 | Thermo Fisher,  Cat#41-4000 |
| NESTIN | Rabbit monoclonal | 1:100 | Abcam  Cat#ab18102 |
| T7 | Rabbit | 1:2000 | Cell Signalling, Cat#13246 |
| GAPDH | Mouse monoclonal | 1:2000 | Abcam  Cat#ab8245 |
| IgG H&L (Alexa Fluor® 488) | Goat anti mouse | 1:200 | Thermo Fisher Scientific  Cat#A11029 |
| IgG H&L (Alexa Fluor® 568) | Goat-anti-rabbit | 1:200 | Thermo Fisher Scientific  Cat#A11036 |
| IgG H&L (Alexa Fluor®488) | Goat-anti-rabbit | 1:200 | Thermo Fisher Scientific  Cat#A11008 |
| IgG H&L (Alexa Fluor® 568) | Goat-anti-mouse | 1:200 | Thermo Fisher Scientific  Cat#A11004 |

**DNA-FISH probes used in this study.**

Dual Labelled Satellite Probe Sets

CytoCell, Cat#LPE 0XYc / LPE 0XYq
