## Supplementary material for "An iPSC-based model of Jacob Syndrome reveals a DNA methylation-independent transcriptional dysregulation shared with X aneuploid cells": Table S9

**Table S9A: TaqMan Gene Expression probes**

| **TaqMan Assays** | **Manufacturer** | **Identifier** |
| --- | --- | --- |
| TaqMan qPCR assay: *TBP* | Thermo Fisher Scientific | Cat#Hs00427620_m1 |
| TaqMan qPCR assay: *OCT4* | Thermo Fisher Scientific | Cat#Hs04260367_gH |
| TaqMan qPCR assay: *NANOG* | Thermo Fisher Scientific | Cat#Hs02387400_g1 |
| TaqMan qPCR assay: *NES* | Thermo Fisher Scientific | Cat#Hs04187831_g1 |
| TaqMan qPCR assay: *TUBB3* | Thermo Fisher Scientific | Cat#Hs00801390_s1 |
| TaqMan qPCR assay: KDM5D | Thermo Fisher Scientific | Cat#Hs00190491_m1 |
| TaqMan qPCR assay: *UTY* | Thermo Fisher Scientific | Cat#Hs01076483_m1 |
| TaqMan qPCR assay: *DDX3Y* | Thermo Fisher Scientific | Cat#Hs00606179_m1 |
| TaqMan qPCR assay: *NLGN4Y* | Thermo Fisher Scientific | Cat#Hs0103434378_s1 |
| TaqMan qPCR assay: *NLGN4X* | Thermo Fisher Scientific | Cat#Hs01934144_s1 |
| TaqMan qPCR assay: *DDX3X* | Thermo Fisher Scientific | Cat#Hs00606179_m1 |
| TaqMan qPCR assay: *UTX* | Thermo Fisher Scientific | Cat#Hs00958902_m1 |
| TaqMan qPCR assay: ZFX | Thermo Fisher Scientific | Cat#Hs01017881_m1 |
| TaqMan qPCR assay: SOX2 | Thermo Fisher Scientific | Cat#Hs01053049_s1 |

**Table S9B: Oligos used for real-time PCR**

| **Oligo Names** | **Manufacturer** | **Exon location** | **Identifier** |
| --- | --- | --- | --- |
| UTX/KDM6A | Sigma-Aldrich | 19-22 | Cat#8821625448-000010/20 |

**Table S9C: Oligos used ZFX cDNA amplification prior subcloning**

| **Oligo Names** | **Manufacturer** | **Sequence (5’-3’)** |
| --- | --- | --- |
| ZFX-NheI-FWD | Sigma | AGCTAGCCATCATTTTGGCAAAG |
| ZFX-XhoI-R | Sigma | ATATCTCGAGAGAATTCTTAGGGCAG |

**Table S9D: siRNAs used for silencing**

| **siRNA Names** | **Manufacturer** | **Sequence** |
| --- | --- | --- |
| hs.Ri.UTY.13.2 | IDT (Integrated DNA technologies) | Not available |
| hs.Ri.ZFY.13.1 | IDT (Integrated DNA technologies) | Not available |
| hs.Ri.DDX3Y.13.3 | IDT (Integrated DNA technologies) | Not available |
| hs.Ri.NLGN4Y.13.2 human | IDT (Integrated DNA technologies) | Not available |
| Human Scrambled Negative Control DsiRNA | IDT (Integrated DNA technologies) | Catalog # 51-01-19-09 |
